# Annelid functional genomics reveal the origins of bilaterian life cycles

**DOI:** 10.1101/2022.02.05.479245

**Authors:** Francisco M. Martín-Zamora, Yan Liang, Kero Guynes, Allan M. Carrillo-Baltodano, Billie E. Davies, Rory D. Donnellan, Yongkai Tan, Giacomo Moggioli, Océane Seudre, Martin Tran, Kate Mortimer, Nicholas M. Luscombe, Andreas Hejnol, Ferdinand Marlétaz, José M. Martín-Durán

## Abstract

Indirect development with an intermediate larva exists in all major animal lineages^1^, making larvae central to most scenarios of animal evolution^2-12^. Yet how larvae evolved remains disputed. Here we show that temporal shifts (i.e., heterochronies) in trunk formation underpin the diversification of larvae and bilaterian life cycles. Combining chromosome-scale genome sequencing in the slow-evolving annelid *Owenia fusiformis*^13^ with transcriptomic and epigenomic profiling during the life cycles of this and two other annelids, we found that trunk development is deferred to pre-metamorphic stages in the feeding larva of *O. fusiformis*, but starts after gastrulation in the non-feeding larva with gradual metamorphosis of *Capitella teleta* and the direct developing embryo of *Dimorphilus gyrociliatus*. Accordingly, the embryos of *O. fusiformis* develop first into an enlarged anterior domain that forms larval tissues and the adult head. Notably, this also occurs in the so-called “head larvae” of other bilaterians^14,15^, with whom *O. fusiformis* larva shows extensive transcriptomic similarities. Together, our findings suggest that the temporal decoupling of head and trunk formation, as maximally observed in “head larvae”, allowed larval evolution in Bilateria, thus diverging from prevailing scenarios that propose either co-option^10,11^ or innovation^12^ of gene regulatory programmes to explain larva and adult origins.

## Main text

Many animal embryos develop into an intermediate, often free-living stage termed larva, which later metamorphoses into the sexually competent adult^1,2^. Larvae are morphologically diverse and display an array of lifestyles, from active predation and drastic metamorphoses (e.g., planktotrophic larvae) to maternal nourishment and a more gradual transition into the adult phase (e.g., lecithotrophic larvae)^1,2^ (Fig. 1a). Given their broad phylogenetic distribution^2^, larvae are central to major scenarios of animal evolution^2-12^. Yet, these dissent on whether larvae are ancestral^2-7^ or secondarily evolved^10,11^ life stages (Fig. 1b). The “intercalation” hypothesis^10,11^ suggests that larval stages were added to animal life cycles multiple times independently, by co-opting genes and genetic programmes originally expressed in the adult. Conversely, the “terminal addition” scenario^2,3,12^ considers that the ancestral bilaterians resembled existing larvae, and thereby adults convergently evolved through the parallel evolution of adult-specific genetic programmes^12^. How these proposed mechanisms of co-option and innovation might translate to our current understanding of the genetic control of animal embryogenesis remains, however, unclear, and thus larval origins–– and their importance to explain animal evolution––are still contentious.

**Figure 1.**
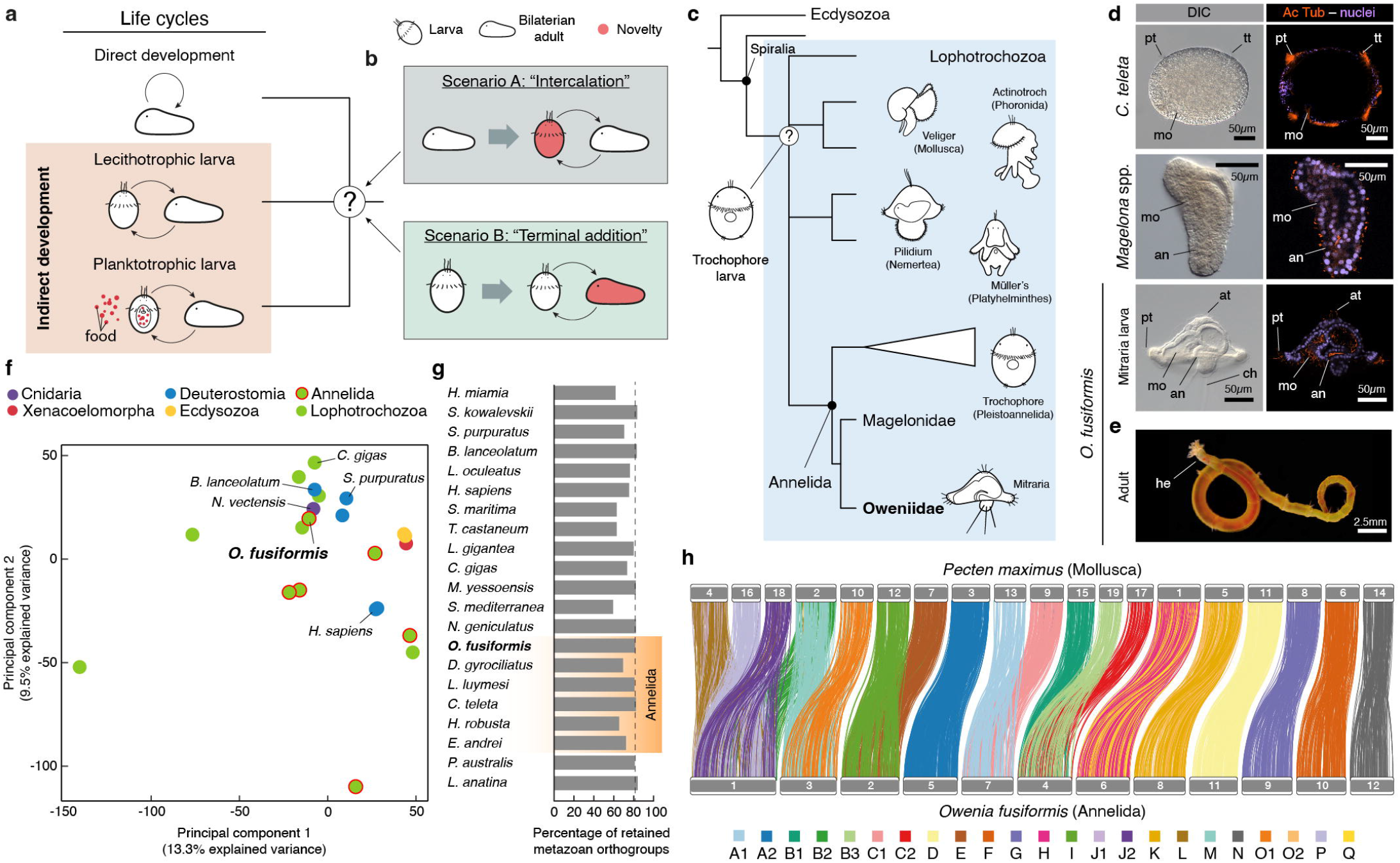
*Owenia fusiformis* has a unique larva and a conservatively evolving genome. **a**, Bilaterian life cycles and larval forms. Within indirect developers, lecithotrophic larvae rely on yolk nutrients during dispersal, while planktotrophic larvae feed on plankton. **b**, Major scenarios for the evolution of bilaterian life cycles and larval forms. The “intercalation” scenario deems bilaterian adults ancestral and larvae secondary specialisations that evolved independently in certain lineages. Opposed to this, the “terminal addition” scenario considers that bilaterian larvae are ancestral and that adult forms evolved secondarily. **c**, A trochophore larval type has been proposed to be ancestral to Spiralia or even Protostomia (Ecdysozoa + Spiralia) and give rise to the diversity of larval forms found in lophotrochozoan taxa. **d**, The larval forms of oweniids and magelonids are unlike other annelid larvae. Differential interface contrast (DIC) images and z-stack confocal laser scanning views of a stage 5 metatrochophore of *C. teleta*, a *Magelona* spp. Larva, and a *O. fusiformis* mitraria stained for DAPI and acetylated α-tubulin. **e**, Image of an adult of *O. fusiformis*. **f**, Principal component analysis of metazoan gene complements demonstrates that *Owenia* clusters with other slow-evolving lineages. See Extended Data Fig. 1g for a fully labelled graph. **g**, Percentage of retained pre-metazoan and metazoan orthogroups per species. Dotted vertical line represents the value for *O. fusiformis*. **h**, Karyotypic correspondence between *O. fusiformis* and *Pecten maximus*, which exemplifies the ancestral spiralian chromosome complement. Each colour represents an ancestral bilaterian linkage group. Schematic drawings are not to scale. At: apical tuft; an: anus; ch: chaetae; he: head; mo: mouth; pt: prototroch; tt: telotroch.

The trochophore^16^ is a widespread larval type generally characterised by an apical sensory organ and a pre-oral locomotive ciliary band^17^ that is typically assigned to Annelida and Mollusca but also potentially to related clades within Lophotrochozoa^18^ (Fig. 1c). Classically exemplified by those of annelid worms, trochophore larvae are pivotal to the “terminal addition” scenario^2,19^, which regards this larval type a vestige of the last common adult ancestor to Protostomia^2,3^, or even Bilateria^20^ (Fig. 1c). Annelids, however, exhibit diverse life cycles and larval morphologies, including species with direct and indirect development and either planktotrophic or lecithotrophic larvae^21^. Notably, the groups Oweniidae and Magelonidae––which form Oweniida, the sister group to all other annelids^13^––exhibit distinctive planktotrophic larvae (Fig. 1c, d). In particular, the oweniid larva, commonly referred to as “mitraria”^22^, has an enlarged pre-oral region and a single bundle of posterior chaetae, as well as a pair of nephridia and a long monociliated ciliary band alike those of phylogenetically distant larvae of echinoderms (e.g., sea urchins) and hemichordates (e.g., acorn worms)^23-25^. Yet, oweniids exhibit many developmental characters considered ancestral to Annelida, and even Lophotrochozoa as a whole^26,27^, as well as similarities in larval molecular patterns with other annelid trochophore and bilaterian larvae^25,26,28,29^. Therefore, Annelida, with its diversity of life cycles and larval forms, but generally conserved early embryogenesis and adult body plans, emerges as an excellent model to investigate how larval traits evolve, and thereby formulate and assess hypotheses on the origin of animal life cycles.

Here, we characterise the reference chromosome-scale genome assembly of the oweniid *Owenia fusiformis* Delle Chiaje, 1844 (Fig. 1e) and perform a comprehensive comparative study of the developmental transcriptomes and regulatory genomes underpinning the formation of its planktotrophic larva, as well as the lecithotrophic larva and direct developing embryo of the annelids *Capitella teleta*^30^ and *Dimorphilus gyrociliatus*^31^, respectively. Diverging from traditional scenarios, our extensive dataset, in comparison with those from other lophotrochozoan and bilaterian taxa, provides compelling evidence that heterochronic shifts in trunk development contribute to the evolution of life cycles in Bilateria.

## O. fusiformis *has a conservatively evolving genome*

To characterise the transcriptomic and genomic regulatory basis for larval development in Annelida, we first generated a chromosome-scale reference assembly for *O. fusiformis* combining PacBio long-reads, 10x Genomics linked-reads, optical mapping, and Hi-C scaffolding (Supplementary Fig. 1a). Consistent with flow cytometry and *k*-mer estimations, the haploid assembly spans 505.8 Mb (Supplementary Fig. 1b–e), exhibiting 12 chromosome-scale scaffolds that encompass 89.2% of the assembly (Supplementary Fig. 1f, g). Almost half of the assembly consists of repeats (43.02%; largely DNA transposons) acquired steadily during evolution (Extended Data Fig. 1a–c; Supplementary Table 1). Using transcriptomic data from 14 developmental stages and 9 adult tissues (Supplementary Fig. 1a), we annotated 26,966 protein-coding genes and 31,903 transcripts, representing a nearly complete (97.5%) set of metazoan BUSCO genes (Supplementary Fig. 1g). Gene family reconstruction and principal component analysis on gene content across 22 animal genomes nested *O. fusiformis* within other non-annelid lophotrochozoan species (Fig. 1f), supporting that *O. fusiformis* has fewer gene family gains and losses, and retains more ancestral metazoan orthogroups than other annelid taxa (Fig. 1g; Extended Data Fig. 1d –g; Supplementary Tables 2–7). Indeed, we identified an ortholog of *chordin*, a bone morphogenetic protein inhibitor involved in dorsoventral patterning across Bilateria^32^ and thought to be lost in annelids^33^ (Extended Data Fig. 2a–f), which is asymmetrically expressed around the blastopore lip of the gastrula and antero-ventral oral ectoderm of the larva in *O. fusiformis* (Extended Data Fig. 2g, h). At a high-order genomic organisation, *O. fusiformis* has globally retained the ancestral bilaterian linkage, exhibiting chromosomal fusions that are present in molluscs and even nemerteans (Fig. 1h; Extended Data Fig. 1h, i), and less lineage-specific chromosomal rearrangements than other annelids (Extended Data Fig. 1h). Therefore, *O. fusiformis* genome contains a more complete gene repertoire and ancestral syntenic chromosomal organisation than those reported for other annelid lineages, which together with its phylogenetic position and conserved early embryogenesis^26,27^ makes it a key lineage to reconstruct the evolution of Annelida and Lophotrochozoa generally.

### Heterochronic shifts in gene regulatory programmes underpin annelid life cycles

To identify transcriptomic changes underpinning distinct life cycles in Annelida, we compared temporal series of embryonic, larval and competent/juvenile transcriptomes of *O. fusiformis* and *C. teleta*, two indirect developers with planktotrophic and lecithotrophic^30^ larvae, respectively (Fig 1d), and *D. gyrociliatus*, a direct developer^31,34^ (Fig. 2a; Supplementary Fig. 2; Supplementary Tables 13–21). In these three species, gene expression increases with time (Fig. 2b; Supplementary Fig. 3) and transcriptional dynamics are overall similar, especially during early embryogenesis (Fig. 2c, d; Supplementary Table 22). Yet, while *C. teleta* and *D. gyrociliatus* show increasing transcriptomic divergence with each other as they develop into adult stages, the maximal transcriptomic divergence between these annelids and *O. fusiformis* occurs with the planktotrophic mitraria (Fig. 2c, d). Soft clustering of all expressed transcripts during development generated an optimal number of 12 distinct clusters of temporally co-regulated genes in *O. fusiformis* and *C. teleta*, and 9 clusters in *D. gyrociliatus* (Extended Data Fig. 3a–c; Supplementary Tables 23–26). In all three species, only one cluster (1,457 transcripts, 4.6% in *O. fusiformis*; 2,206 transcripts, 5.54% in *C. teleta*; and 1,847 transcripts, 10.75% in *D. gyrociliatus*) showed a bimodal activation involving early embryonic stages (blastulae in *O. fusiformis* and *C. teleta*; early embryogenesis in *D. gyrociliatus*) and competent/juvenile/adult stages (Extended Data Fig. 3a –c). These clusters are enriched in gene ontology (GO) terms associated with core cellular processes, such as catabolism and cellular transport (in *O. fusiformis*), oxidative phosphorylation (OXPHOS) metabolism (in *C. teleta*) and RNA processing, metabolism, and catabolism (in *D. gyrociliatus*) (Extended Data Fig. 3d, e, Supplementary Fig. 4–11, Supplementary Tables 27–31). Indeed, translation and metabolism predominate in clusters of early developmental phases in the three annelids, while cell communication and signalling, morpho- and organogenesis are enriched in later stages of development (Extended Data Fig. 3d). Thus, regardless of their life cycle, annelids share global transcriptional dynamics during development, yet adults and the planktotrophic larva are the most transcriptionally distinct stages.

Divergence in larval developmental programmes (null expectation; Fig. 2e) or alternatively, changes in expression levels of otherwise conserved larval transcriptomes and temporal shifts (i.e., heterochronies), could explain transcriptomic differences between annelid life cycles (Fig. 2e). To test these, we performed pairwise inter-species comparisons of gene (Extended Data Fig. 4a, b) and transcription factor (Fig. 2f; Supplementary Fig. 3, Supplementary Tables 32, 33) composition among clusters of temporally co-regulated genes. In agreement with transcriptional similarity comparisons (Fig. 2c, d), early, followed by late clusters—pre- and post-larval in indirect developers, respectively (Extended Data Fig. 4c)—are the most conserved phases in the three comparisons when all genes are considered (Extended Data Fig. 4a –d). However, transcription factors that are deployed post-larva in indirect development are consistently shifted to early embryogenesis in direct development (Fig. 2f, dotted rectangles; Extended Data Fig. 4e), including 28 transcription factors common to *O. fusiformis* and *C. teleta* and that are involved in a variety of processes (Fig. 2g; Supplementary Tables 34, 35), from nervous system (e.g., *pax6*^35^) and mesoderm (e.g., *foxF*^29^) development to axial patterning (e.g., *Hox1* and *Hox4*^36^). Notably, the overall expression of these 28 genes is also temporally shifted between indirect developing annelids, with the maximum level of expression occurring earlier in *C. teleta* than in *O. fusiformis* (Fig. 2g). Consistently, 2,583 genes also exhibit temporal shifts between the larvae of *O. fusiformis* and *C. teleta* (Extended Data Fig. 4f). These include 105 transcription factors, but most are enzymes and structural genes (Extended Data Fig. 4f –g, Supplementary Fig. 12, 13, Supplementary Tables 36–42) that likely reflect the different biology of these two larvae. For example, genes involved in chitin biosynthesis are among those expressed early in *O. fusiformis* but late in *C. teleta* (Supplementary Fig. 14, Supplementary Tables 43, 44), in agreement with the conspicuous chaetae of the mitraria larva that are absent in the capitellid larva (Fig. 1d). Conversely, the autophagy pathway, which is involved in yolk catabolism^37^, is deployed earlier in the lecithotrophic larva of *C. teleta* (Extended Data Fig. 4i, Supplementary Tables 45, 46). Therefore, although it is likely a combination of factors that explain transcriptional differences, heterochronic changes—sometimes involving just a few key genes related to arange of genetic programmes—correlate, and might account for, life cycle and larval differences in Annelida.

**Figure 2.**
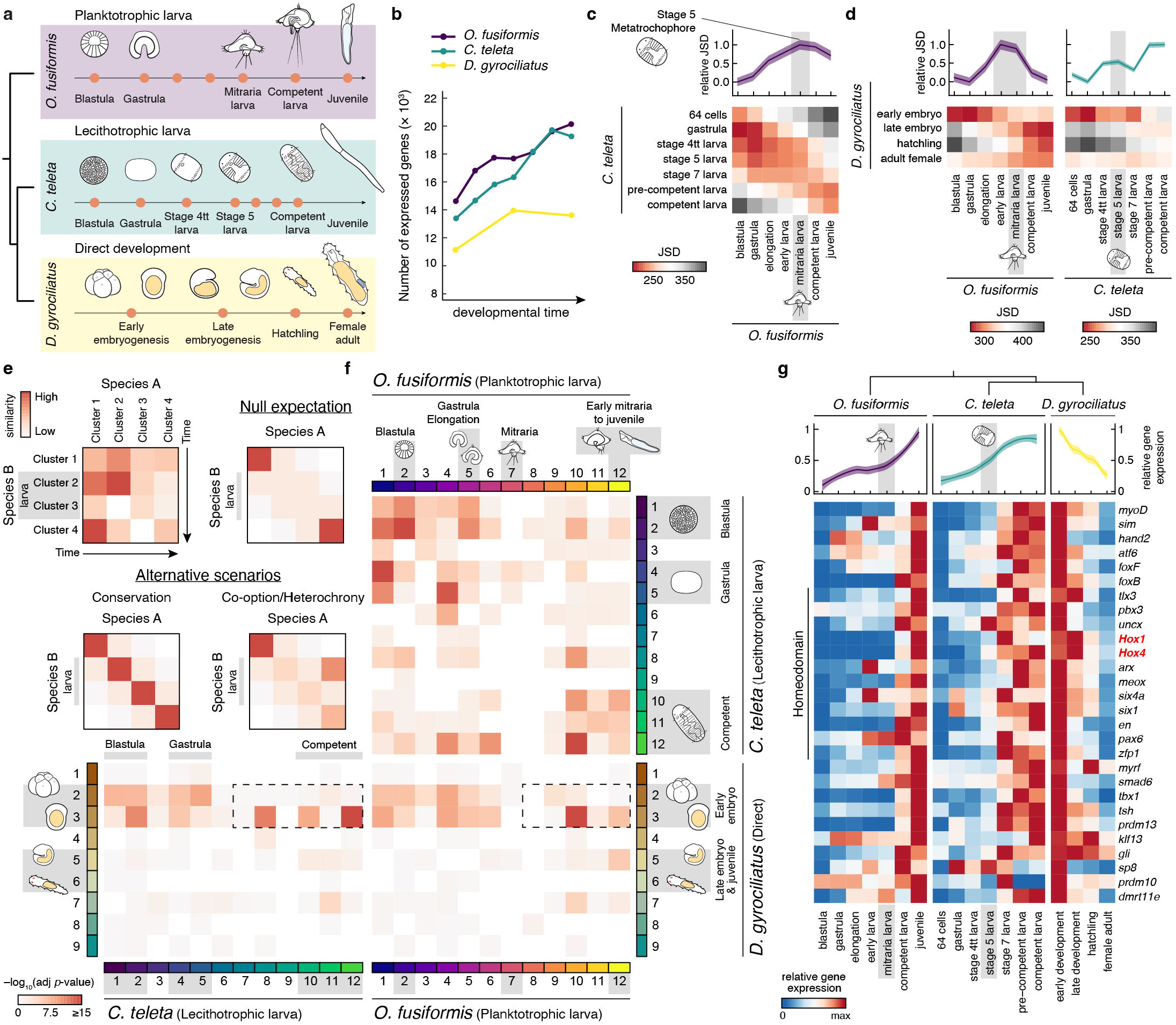
Heterochronic shifts in gene regulatory programmes underpin annelid life cycle diversification. **a**, Experimental design of the comparative developmental RNA-seq time courses. Orange circles highlight stages of *O. fusiformis, C. teleta*, and *D. gyrociliatus* development sampled for bulk RNA-seq. **b**, Number of expressed genes during annelid development. See Supplementary Fig. 3c for fully labelled plots. **c, d**, Heatmaps of pairwise transcriptomic Jensen–Shannon Divergence (JSD) between *O. fusiformis* and *C. teleta* (**c**), and between *D. gyrociliatus* and either *O. fusiformis* (**d**, left) or *C. teleta* (**d**, right). Relative JSD of the *C. teleta* or *O. fusiformis* stages of minimal divergence to each corresponding stage is shown on top. **e**, Schematics of the orthogroup overlap strategy. Chronologically ordered clusters of co-expressed genes from two distinct species with similar developmental gene expression dynamics would render a similarity diagonal (null expectation). Alternative scenarios include either lack of overlap in divergence or innovation situations, or timing differences when co-opted or temporally shifted genetic programmes are present. **f**, Similarity heatmaps showcasing the orthogroup overlap between the transcription factors contained in clusters of co-regulated genes obtained by soft *k*-means clustering, between all three studied annelid taxa. Time points associated to key clusters are shown for all three species. Dotted black lines encompass the sharp timing expression differences of a significant number of transcription factors shifted from post-larval expression in indirect developers to early embryogenesis in *D. gyrociliatus* can be observed. **g**, Heatmaps of gene-wise expression (bottom) and average expression dynamics (top) during *O. fusiformis, C. teleta*, and *D. gyrociliatus* development of the set of 28 single copy ortholog transcription factors shifted from late expression in both *O. fusiformis* and *C. teleta* to early expression in *D. gyrociliatus*. Up to 43 % (12) of these belong to the homeodomain class of transcription factors.

### *Trunk development is delayed to pre-metamorphosis in* O. fusiformis

Homeodomain transcription factors control many, often conserved developmental processes^38^ and are the largest class among the 28 transcription factors exhibiting temporal shifts between annelids with different life cycles (Fig. 2g). Indeed, homeodomain genes are enriched in the competent larva in *O. fusiformis* but are prevalent among genes active from stage 5 larva onwards in *C. teleta* (Extended Data Fig. 4j). Accordingly, *Hox* genes—a conserved family involved in anterior-posterior trunk regionalisation in Bilateria^39^—are among the most upregulated genes in the competent mitraria larva (Extended Data Fig. 5b). Similar to *C. teleta*^36^ and *D. gyrociliatus*^34^ (although the latter lacks *Lox2* and *Post1*), *O. fusiformis* has a conserved complement of 11 *Hox* orthologues arranged as a compact, ordered cluster in chromosome 1, except for *Post1*, which is located downstream on that same chromosome (Extended Data Fig. 5a, c, d; Supplementary Table 47). However, while *C. teleta* and *D. gyrociliatus* deploy *Hox* genes during or soon after gastrulation^34,36^ (Extended Data Fig. 5e), *O. fusiformis* does not express *Hox* genes during embryogenesis to pattern the larval body (Fig. 3a; Extended Data Fig. 5e, f). Instead, *Hox* genes are expressed in the trunk rudiment during larval growth, already in an anterior–posterior staggered pattern that is retained in the juvenile after metamorphosis (Fig. 3a –c; Extended Data Fig. 5f). This late activation of *Hox* genes is not unique to *O. fusiformis*, but also occurs in the planktotrophic trochophore of the echiuran annelid *Urechis unicinctus* (Extended Data Fig. 5e; Supplementary Table 48). Therefore, the spatially collinear *Hox* code along the trunk is established at different developmental stages depending on the life cycle mode in Annelida.

**Figure 3.**
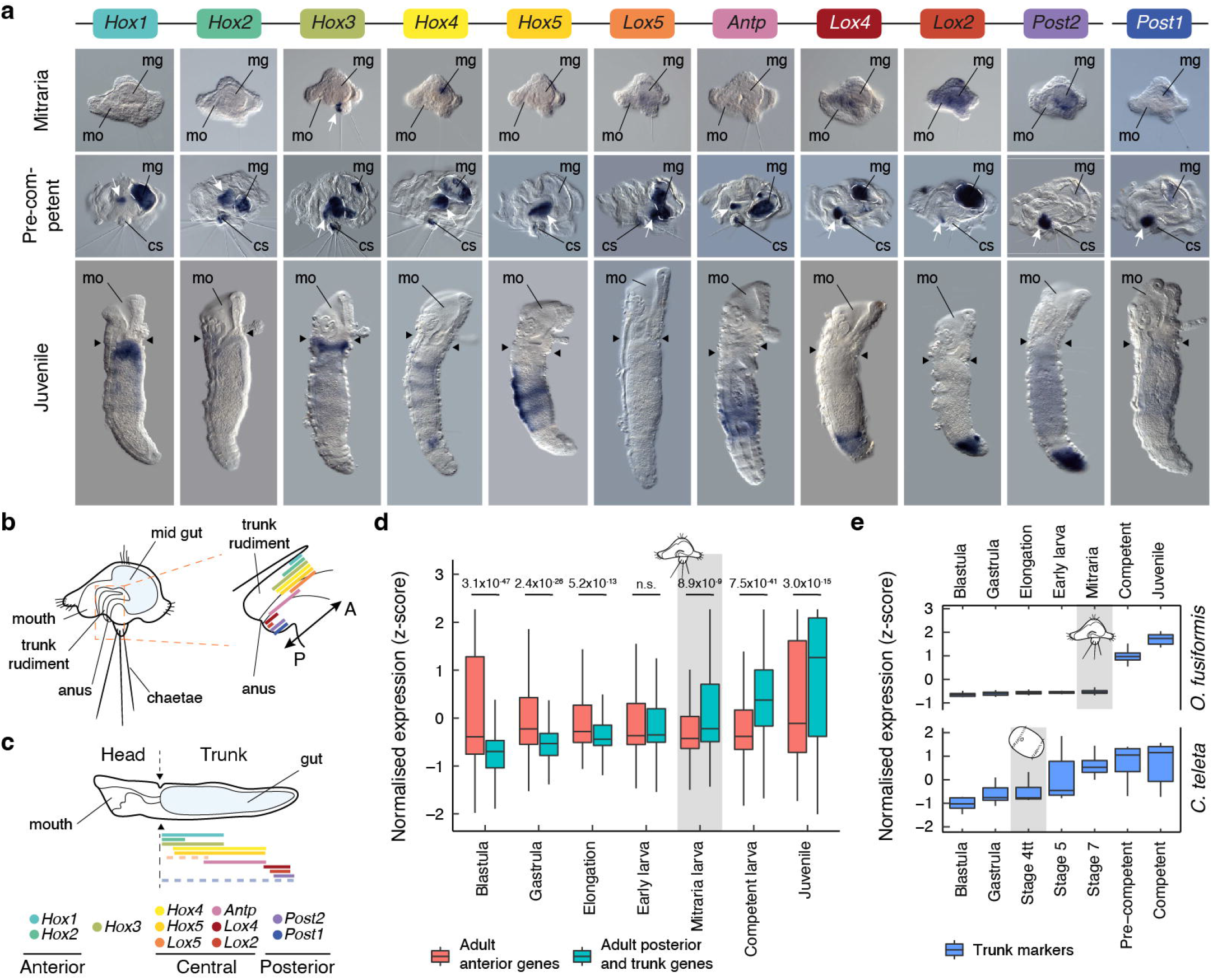
Trunk development is delayed to pre-metamorphosis in *O. fusiformis*. **a**, Whole mount *in situ* hybridisation of *Hox* genes at the mitraria, pre-competent, and juvenile stages of *O. fusiformis*. Only *Hox3* appears to be expressed at the mitraria stage (white arrow). *Hox* genes show spatial collinearity along the anterior-posterior axis at the developing trunk (white arrows in the pre-competent larva) and juvenile. Dotted lines in the competent larva panels indicate background from the gut content. Black arrowheads in the juvenile panels indicate head to trunk boundary. cs: chaetal sack; mg: mid gut; mo: mouth. **b, c** Schematic representations of the expression of *Hox* genes in the trunk rudiment of the competent larva (**b**) and juvenile trunk (**c**). A: anterior; P: posterior. Drawings are not to scale, and schematic expression domains are approximate. **d**, Expression dynamics of anterior and posterior and trunk genes expressed in corresponding adult tissues during *O. fusiformis* development. *P-*values were derived from two-tailed Student’s *t*-tests and adjusted with the Bonferroni method for multiple testing correction. n.s.: not significant. **e**, Expression dynamics of *in situ* hybridisation-validated trunk markers throughout *O. fusiformis* (top) and *C. teleta* (bottom) development. In contrast with the gradual onset from the stage 4tt larva stage in *C. teleta*, trunk markers deploy markedly but only after the mitraria larva stage in *O. fusiformis*.

To determine whether the different timings of trunk patterning is limited to *Hox* genes or involve broader genetic programmes associated with trunk development in annelids, we used tissue-specific adult transcriptomes to define a set of 1,655 anterior and 407 posterior/trunk genes in *O. fusiformis* (Extended Data Fig. 6a –c; Supplementary Tables 49–53). Anterior genes are significantly more expressed during embryogenesis (Fig. 3d; Extended Data Fig. 6d, e). On the contrary, posterior/trunk genes are upregulated at the mitraria stage and significantly outweigh the expression dynamics of anterior genes from that stage on (Fig. 3d; Extended Data Fig. 6d, f). Indeed, anterior, trunk, and posterior genes for which there is spatially resolved expression data follow different temporal dynamics in *O. fusiformis, C. teleta*, and *D. gyrociliatus* (Extended Data Fig. 6g –l; Supplementary Tables 54–56). In *O. fusiformis*, trunk (e.g., *nkx6, msx* paralogs^28^) and posterior genes (e.g., *evx*^29^, *wnt1*^27^) concentrate in a small ventral area and around the anal opening of the larva^27-29^ and increase in spatial range and expression level as the trunk forms. On the other hand, anterior genes (e.g., *foxQ2* genes^40^, *gsc*^29^, *otx*^29^) pattern most of the mitraria and their expression remain fairly stable during development (Fig. 3d; Extended Data Fig. 6g, h). In contrast, posterior and anterior genes follow similar dynamics in *C. teleta* (Extended Data Fig. 6i, j) and trunk genes upregulate already post-gastrula in both *C. teleta* and *D. gyrociliatus* (Fig. 3e; Extended Data Fig. 6i –l). Therefore, trunk development, which initially occurs from lateral growth of the trunk rudiment^22,30,41^, is deferred to pre-metamorphic stages in planktotrophic annelid trochophores^14,42^ compared to annelids with lecithotrophic larvae^36,43^ and direct developers^34^.

### Chromatin dynamics support heterochronic shifts in Hox regulation between annelid larvae

To investigate the genomic regulatory basis for the temporal shift in trunk development between annelid larvae, we profiled open chromatin regions with ATAC-seq at five equivalent developmental stages based on cross-species minimal transcriptomic distance in *O. fusiformis* and *C. teleta* (Fig. 4a; Supplementary Fig. 15; Supplementary Tables 57–60). In total, we identified 63,726 and 44,368 consensus regulatory regions in *O. fusiformis* and *C. teleta*, respectively, mostly abundant within gene bodies (69.69% and 51.53%, respectively; from transcription start to end sites) rather than in promoters (12.92% and 20.53%) and distal intergenic regions (17.38% and 27.93%) (Extended Data Fig. 7a, b; Supplementary Fig. 16). The largest changes in peak accessibility occur in the mitraria (in *O. fusiformis*) and stage 5 larva (in *C. teleta*) (Supplementary Fig. 16; Supplementary Tables 61–66), with a general increase in promoter peaks (in *O. fusiformis*) and distant intergenic regulatory elements (in both species) during development (Fig. 4b). Soft clustering revealed, however, that most regulatory regions act before the start of trunk formation in *O. fusiformis* (clusters 1 to 9; 47,652 peaks; 76. 72%), while the number of accessible regions with a maximum of accessibility before (clusters 1 to 7; 23,678 peaks; 51.45%) and after (clusters 8 to 12; 22,341; 48.56%) the onset of trunk development (stage 4tt larva) are comparable in *C. teleta* (Fig. 4c; Supplementary Tables 67–69). Moreover, regulation of genes involved in morpho- and organogenesis, as well as neurogenesis, concentrates in late clusters in *O. fusiformis*, but unfolds more continuously in *C. teleta* (Supplementary Fig. 17–21). Therefore, different dynamics of temporally co-regulated accessible chromatin regions occur during development and larva formation in *O. fusiformis* and *C. teleta*.

**Figure 4.**
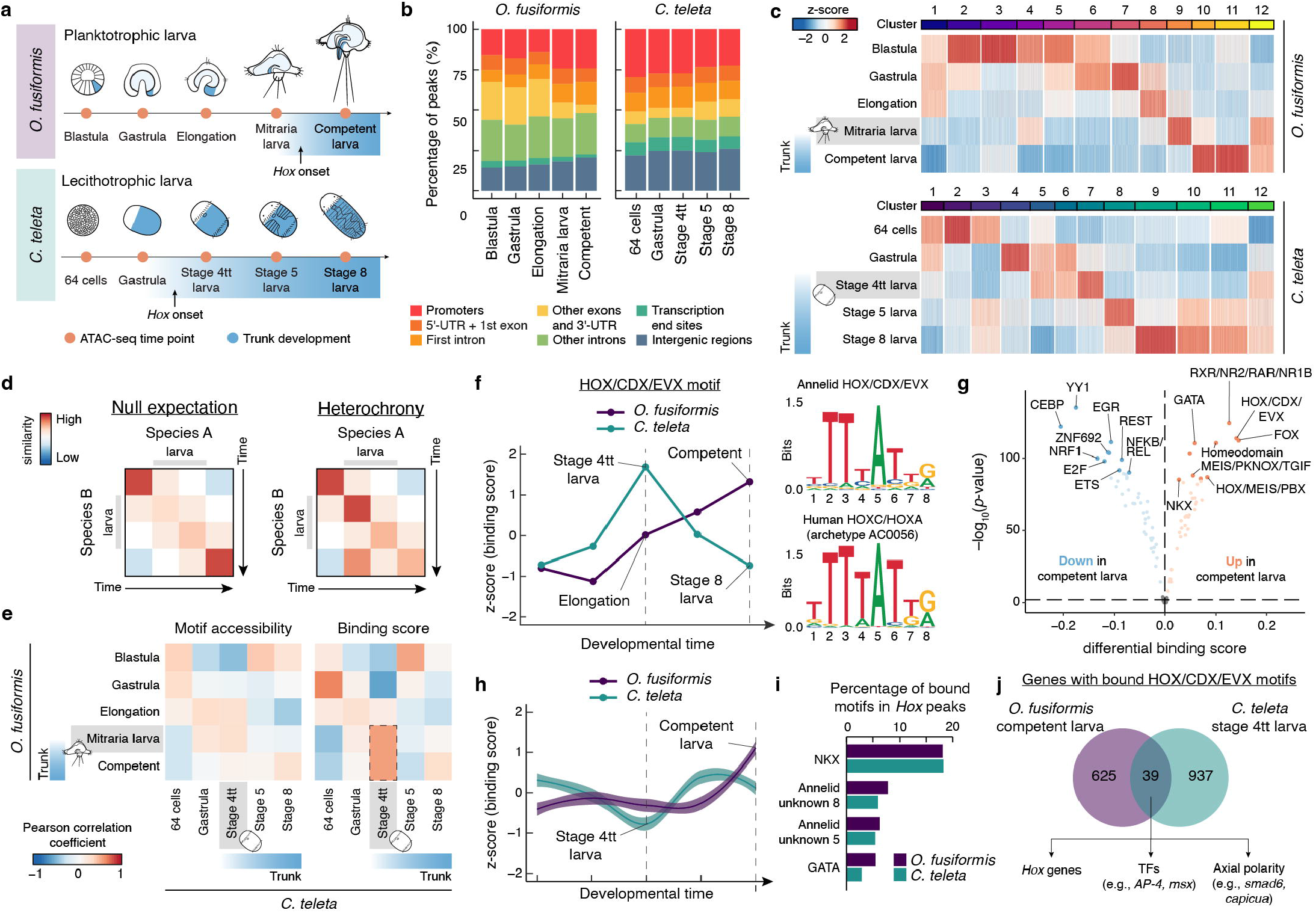
Chromatin dynamics support heterochronic shifts in *Hox* regulation between annelid larvae. **a**, Experimental design of the comparative developmental ATAC-seq time courses, highlighting with orange circles the stages of *O. fusiformis* and *C. teleta* development sampled for bulk ATAC-seq. *Hox* expression onset, trunk development, and larval trunk domains are indicated in blue. **b**, Stacked bar plots showing the proportion of called peaks per developmental stage classified by genomic feature for *O. fusiformis* (left) and *C. teleta* (right). **c**, Heatmap of normalised peak accessibility of the soft clustered consensus ATAC-seq peak sets of *O. fusiformis* (top) and *C. teleta* (bottom). **d**, Schematics of the cross-species ATAC-seq comparison strategy. Our null expectation entails lack of similarity between regulatory dynamics at larval stages. Alternatively, heterochronic shifts of regulatory programmes could support the heterochronic shifts in gene expression observed between *O. fusiformis* and *C. teleta*. **e**, Correlation matrices of motif archetype accessibility and transcription factor binding score (TFBS). Dotted black lines highlight the high TFBS correlation and subsequent heterochronic shift between the mitraria and competent larva of *O. fusiformis* and the stage 4tt larva of *C. teleta*. **f**, TFBS dynamics for the HOX/CDX/EVX motif archetype during *O. fusiformis* (purple) and *C. teleta* (blue) development. Sequence logo of the annelid archetype (top) shows the substantial similarity to the human homolog (bottom). **g**, Volcano plot of differential transcription factor binding between the mitraria larva and the competent larva stages of *O. fusiformis*. **h**, Average TFBS dynamics of all motifs in the peaks of the *Hox* genes clusters. **i**, Most abundant bound motifs in peaks of the *Hox* genes clusters. **j**, Downstream regulated genes by transcription factors bound to the HOX/CDX/EVX motif archetype.

Divergence in the regulatory programmes controlling larva development (null expectation), or alternatively, temporal changes in the deployment of common trunk regulatory programmes might explain the heterochronic shifts in trunk development in annelid larvae (Fig. 4d). To assess these scenarios, we predicted transcription factor-binding motifs on ATAC-seq peaks in *O. fusiformis* and *C. teleta*, which we clustered into 95 and 91 consensus motifs, respectively (Supplementary Fig. 22; Supplementary Tables 72–74). Among these, 51 are common to both species, including 33 that are robustly assigned to a known transcription factor class based on sequence similarity to human motif archetypes (Supplementary Tables 74–76). Motif accessibility and binding—as proxies of regulatory activity^44^—are well self- and cross-species correlated in both species (Extended Data Fig. 7a –c; Supplementary Tables 77–80) and reveal clusters of co-regulatory activity during *O. fusiformis* and *C. teleta* development (Supplementary Fig. 23, 24). The comparison of accessibility and binding dynamics of the 33 common annotated motifs between *O. fusiformis* and *C. teleta* show a temporal shift of 4tt larva of *C. teleta* (Fig. 4e, dotted rectangle; Supplementary Fig. 25–27). Seven motifs (21%) followed this pattern (Extended Data Fig. 7f), including one with high similarity to the human HOX/CDX/EVX motif archetype (Fig. 4f; Supplementary Fig. 27) and that is among the most upregulated and overrepresented motifs at the competent stage in *O. fusiformis* (Fig. 4g; Extended Data Fig. 7g; Supplementary Fig. 27). Indeed, motif binding in *Hox* regulatory elements support a change of global regulation of the *Hox* cluster at the competent and stage 4tt in *O. fusiformis* and *C. teleta*, respectively (Fig. 4h; Supplementary Fig. 29), largely mirroring the transcriptional onset of these genes and the start of trunk development in the two species^36^ (Extended Data Fig. 5e). Motifs assigned to NKX and GATA factors, as well as two annelid-specific motifs of unclear assignment, are the most abundant bound motifs in the *Hox* cluster in both species (Fig. 4i; Supplementary Tables 81, 82). Notably, *nkx* and *gata* genes are expressed in the juvenile rudiment^28^ and along the annelid trunk^28,45^ in *O. fusiformis* and *C. teleta*, and the putative GATA motif follows similar binding dynamics in these species than the HOX/CDX/EVX motif archetype (Extended Data Fig. 7c; Supplementary Fig. 27). However, only 39 one-to-one orthologs with bound HOX/CDX/EVX motifs are common to *O. fusiformis* and *C. teleta* at the time of maximum motif binding, albeit those include *Hox* themselves and a reduced set of transcription factors—some involved in trunk development, such as *msx*^28^—and axial polarity genes (Fig. 4j; Supplementary Tables 83, 84). Together, our epigenomic data reinforces that different regulatory dynamics of the *Hox* cluster—possibly by a reduced common set of upstream regulators––result in temporal shifts of the binding activity of *Hox* genes and expression of trunk patterning genes, likely accounting for the developmental and morphological differences between planktotrophic and lecithotrophic annelid larvae.

### The larva of Owenia does not rely on novel genes

Novel genes account for a significant proportion of some trochophore and other larval transcriptomes^7,46^, as they might be associated with the development of larval-specific characters (e.g., ciliary bands)^7,46,47^. Therefore, recruitment of novel genes at larval stages could also explain transcriptomic—and morphological—differences at this stage (Fig. 2c, d; Extended Data Fig. 4a, b). To define the contribution of novel genes at each developmental stage in annelids with different life cycles, we classified all predicted transcripts in seven different phylostrata according to their time of origin (Extended Data Fig. 8a; Supplementary Tables 85–87). Genes of metazoan and pre-metazoan origin (phylostratum 1) tend to peak, dominate and be enriched at earlier developmental stages in the three species (Fig. 5a; Extended Data 8h–j), whereas younger genes are more highly expressed in competent and juvenile stages (Fig. 5a, Extended Data Fig. 8b –g). Notably, species-specific genes are more expressed and enriched in the juvenile stages of *O. fusiformis* and *D. gyrociliatus*, but in the blastula and gastrula of *C. teleta* (and to some extent also at the blastula stage in *O. fusiformis*; Fig. 5a). Indeed, species-specific genes follow lineage-specific dynamics (Supplementary Fig. 30). At larval stages, however, species-specific genes are only enriched in *C. teleta*, while the mitraria shows enrichment of genes of metazoan and annelid origin (Extended Data Fig. 8h, i). Therefore, genes of different evolutionary origins contribute to the development of annelid larvae, suggesting that the increased use of novel genes in some lophotrochozoan larvae^7,46^ might be due to lineage-specific traits, such as the shell primordium of molluscan trochophores^7^ and perhaps even ciliary bands with multiciliated cells^48^, which are absent in oweniid larvae.

**Figure 5.**
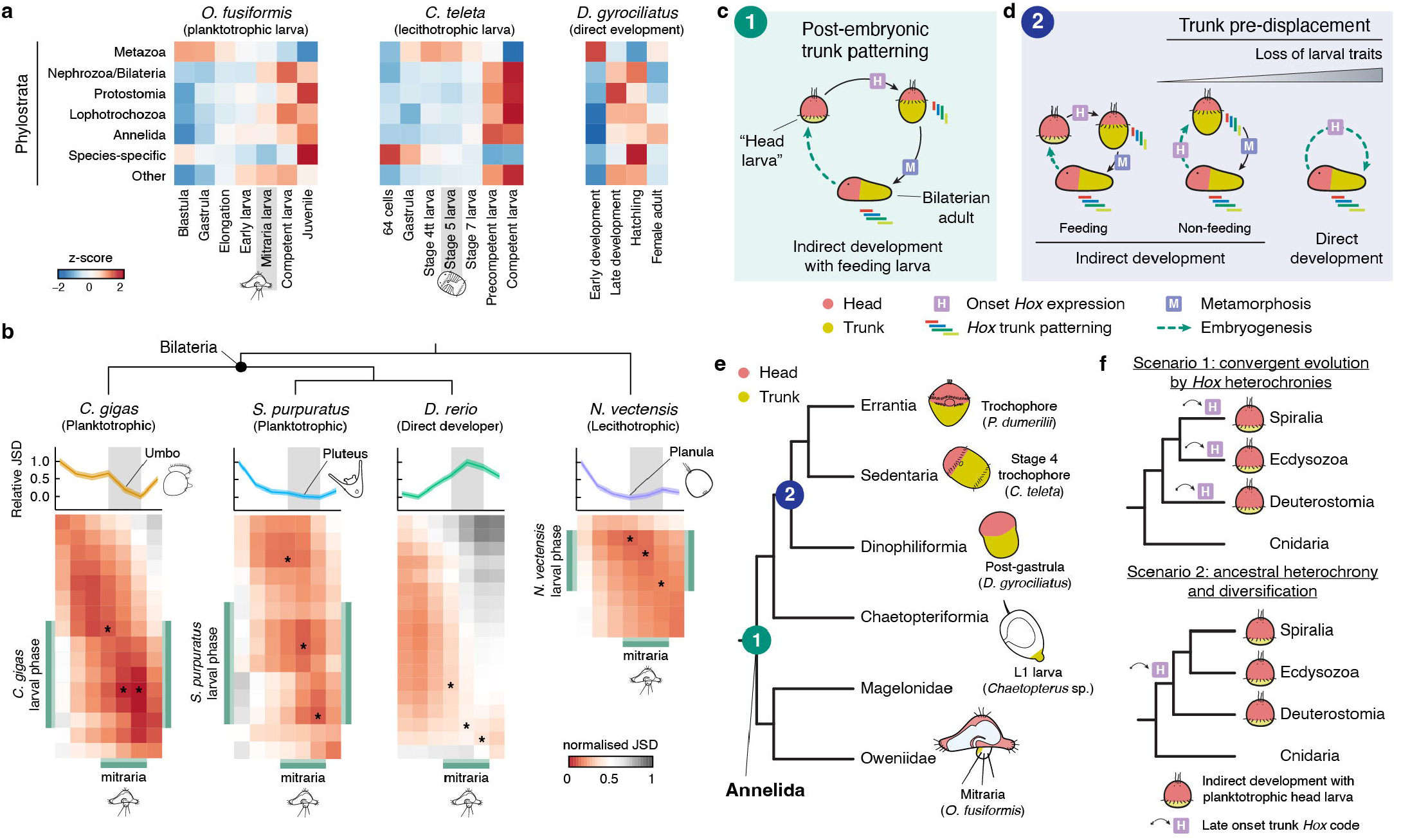
Gene novelties and the evolution of life cycles in Annelida and Bilateria. **a**, Expression dynamics of each phylostratum by developmental stage in all three annelids, calculated from the 75% percentile of a quantile-normalised matrix of gene expression levels. **b**, Heatmaps of pairwise normalised Jensen-Shannon divergence (JSD) between *O. fusiformis* and *C. gigas, S. purpuratus, D. rerio* and *N. vectensis*. Asterisks indicate the stages of minimal JSD of each species to the larval phase of *O. fusiformis*. Larval phases are highlighted in green. On top, relative JSD from stages of minimal divergence to each *O. fusiformis* developmental stage. Confidence intervals represent the standard deviation from 250 bootstrap resamplings of the single copy ortholog set. **c**, Schematic drawing of the life cycle and patterning events in a bilaterian with indirect development with a feeding larva like the annelid *O. fusiformis*. Embryogenesis results in a larva with mostly anterior ectodermal fates from which larval organs and the adult head forms. The onset of trunk differentiation and *Hox* gene expression occurs later, with larval growth pre-metamorphosis. **d**, Schematic drawings of the three main types of life cycles and the timing of *Hox* gene expression in bilaterians. Compared to indirect development with feeding larvae, lineages with non-feeding larvae and direct development pre-displace (i.e., initiate earlier) trunk differentiation and *Hox* gene expression. Larval organs are reduced in non-feeding larvae and absent in direct development. **e**, Proposed evolutionary scenario for larval and life cycle evolution in Annelida. Post-embryonic trunk patterning is likely an ancestral condition (1) with the convergent pre-displacement of trunk differentiation to embryogenesis (2) concurring with the evolution of indirect development with feeding larva and direct development. Drawings are not to scale. **f**, Alternative scenarios for the evolution of “head larvae” in Bilateria.

### Bilaterian planktotrophic larvae share maximal transcriptional similarity

To assess whether similar transcriptional dynamics to those found in annelids are observed across different life cycles in Metazoa, we extended our comparative transcriptomic approach to nine other animal lineages using single copy one-to-one orthologs and reduced sets of either conserved cross-species single copy orthologs or one-to-one transcription factors (Extended Data Fig. 9a, b; Extended Data Fig. 10a; Supplementary Tables 88–90). In relative terms, global transcriptional dynamics between *O. fusiformis* and other major animal groups tend to be more dissimilar at early development, increasing in similarity as embryos proceed towards juvenile and adult stages (Fig. 5b; Extended Data Fig. 9a, b). The exception is the direct developer *Danio rerio*, for which the mitraria larva is the most dissimilar stage (Fig. 5b), as also occurs when comparing *O. fusiformis* with the direct developing annelid *D. gyrociliatus* (Fig. 2d). However, *O. fusiformis* shares maximal transcriptomic similarities during larval phases with bilaterian species with planktotrophic ciliated larvae and even the planula stages of the cnidarians *Nematostella vectensis* and *Clytia hemisphaerica* (Fig. 5b; Extended Data Fig. 9a –e; Extended Data Fig. 10a). The expression of genes involved in core cellular processes, such as transcription, translation, OXPHOS metabolism, and the assembly of cellular components and complexes, directly contribute to these similarities, likely reflecting common structural and ecological needs of metazoan larvae (Extended Data Fig. 9f, g; Supplementary Tables 91, 92). Yet, regulatory programmes as inferred from similarities in transcription factor expression levels are also maximally similar between those species at larval phases (Extended Data Fig. 9a, b, e). Together, these findings suggest that adult development is generally more similar^10^ than early embryogenesis across major animal lineages, maybe because adult bodies operate on many similar cell biological functions. Additionally, these results also reveal unexpected genome-wide transcriptional—and potentially regulative—similarities between phylogenetically distant animal larvae.

## Discussion

In this study, we report the chromosome-scale genome assembly of the oweniid *O. fusiformis* and extensive gene expression and genomic regulatory profiling during embryonic and larval development in three annelids with distinct life cycles. Together, these datasets provide an unprecedented perspective on life cycle evolution in Bilateria. We show that the planktotrophic larva of *O. fusiformis* defers trunk differentiation to late pre-metamorphic stages and largely develops from anterior ectodermal domains (Fig. 5c). This occurs in other feeding annelid larvae^42^ (Extended Data Fig. 5f), and likely in Chaetopteriformia^49-52^ too, and thus the late differentiation of the adult trunk might be an ancestral trait to Annelida (Fig. 5e). Notably, delaying trunk development to post-larval stages also occurs in phylogenetically distant clades within Lophotrochozoa (e.g., nemerteans^53^ and phoronids^54^), Ecdysozoa (pancrustaceans^15^ and pycnogonids^55^), and Deuterostomia (e.g., echinoderms^56,57^ and hemichordates^58^), whose larvae are generally referred to as “head larvae”^14,15^. By contrast, non-feeding larvae^36,59^ and direct developers^34^ in both Annelida and other bilaterian taxa^60,61^ start to pattern their trunks with or straight after the onset of anterior/head patterning (Fig. 5d, e; Extended Data Fig. 10b), which always takes place before gastrulation in bilaterians^62,63^. Therefore, heterochronic shifts in trunk development correlate with, and possibly account for, the evolution of different life cycles in animals. This mechanism thus differs from predictions that propose either co-option of adult genes into larval-specific gene regulatory programmes^10,11^ (in line with the “intercalation” scenario) or the independent evolution of adult gene regulatory modules^2,3,19^ (as it would be expected in the “terminal addition” scenario) as major drivers of larva and adult evolution, respectively.

The post-embryonic activation of genetic programmes involved in trunk differentiation observed in bilaterian “head larvae” could be lineage-specific innovations associated with the evolution of maximal indirect development^14,15,24,54^. Consequently, “head larvae” evolved convergently by repeatedly delaying trunk differentiation and *Hox* patterning (Fig. 5f). The observed similarities in larval molecular patterns^6,54,58,64^ thus reflect ancient gene regulatory modules that were independently co-opted to develop analogous cell types and larval organs (e.g., apical organs and ciliary bands). In contrast, an ancestral post-embryonic onset of trunk differentiation and *Hox* expression might be the most parsimonious state for Bilateria, based on our current understanding of the timing of trunk-associated expression of *Hox* genes in major bilaterian clades and Cnidaria (Extended Data Figure 10c; Supplementary Fig. 31; Supplementary Tables 93, 94). This ancestral temporal decoupling between head and trunk genetic programmes could thus have facilitated the evolution of larvae, which would then originally share the use of anterior genetic modules for their development (Fig. 5f).

Regardless of the scenario, however, and despite the limitations that whole-embryo transcriptomic and epigenomic datasets without spatial resolution have, our study highlights the importance of heterochronic changes in the deployment of ancient genetic programmes for the diversification of bilaterian life cycles, uncovering a reduced set of candidate genes (Fig. 2g) and regulatory motifs (Extended Data Fig. 7f) that might influence life cycle differences in Annelida, and perhaps even Bilateria. In the future, comparative functional studies of these and other genes are needed to thoroughly dissect the regulatory principles underlying head and trunk development and decipher how temporal changes in gene expression and regulation have shaped the evolution of larval and adult forms in Bilateria.

## Methods

### Adult culture, spawning and in vitro fertilisation

Sexually mature *O. fusiformis* adults were collected from subtidal waters near the Station Biologique de Roscoff and cultured in the lab as described before^26^. *In vitro* fertilisations and collections of embryonic and larval stages were performed as previously described^26^. *Capitella teleta* Blake, Grassle & Eckelbarger, 2009 was cultured, grown, and sifted, and its embryos and larvae were collected following established protocols^30^. *Magelona* spp. were collected in muddy sand from the intertidal of Berwick-upon-Tweed, Northumberland, NE England (∼55.766781, -1.984587) and kept initially in aquaria at the National Museum Cardiff before their transfer to Queen Mary University of London, where they were kept in aquaria with artificial sea water.

### Genome size measurements

To estimate the haploid DNA nuclear content of *O. fusiformis*, we used a flow cytometer Partex CyFlow Space fitted with a Cobalt Samba green laser (532 nm, 100 mW) as described for the annelid *Dimorphilus gyrociliatus*^26^ and adult individuals of *Drosophila melanogaster* as reference. Additionally, we used Jellyfish v.2.3^65^ to count and generate a 31-mer histogram from adaptor-cleaned, short-read Illumina reads (see section below), and GenomeScope 2.0^66^ to obtain an in-silico estimation of the genome size and heterozygosity of *O. fusiformis*.

### Genome sequencing, assembly, and quality check

Ultra-high molecular weight (UHMW) genomic DNA (gDNA) was extracted following the Bionano genomics IrysPrep agar-based, animal tissue protocol using sperm from a single *O. fusiformis* male. UHMW gDNA was cleaned up using a salt:chloroform wash following PacBio’s recommendations before long-read sequencing using PacBio v3.0 chemistry at the University of California Berkeley. A total of 16 SMRT cells of PacBio Sequel were used for sequencing with 600 min movie time, producing a total of 170.07 Gb of data (10.72 million reads, N50 read length between 25.75 kb and 30.75 kb). In addition, we used UHMW gDNA of that same individual to generate a 10x Genomics linked reads library, which we sequenced in an Illumina HiSeq4000 at Okinawa Institute of Science and Technology (OIST) to produce 28.62 Gb of data (141.66 million read pairs). PacBio reads were assembled with CANU v.8.3rc2^67^ assuming ‘batOptions=“-dg 3 -db 3 -dr 1 -ca 500 -cp 50’ and ‘correctedErrorRate=0.065’. Pacbio reads were remapped using pbalign v.0.3.2 and the assembly polished once using Arrow (genomicconsensus, v2.3.2). Then Illumina paired end reads generated with the 10x Genomics linked reads were extracted, remapped using bwa mem v.0.7.17^68^ and used for polishing with Racon v.1.16^69^. Bionano Genomics optical mapping data was used to scaffold the PacBio-based assembly, which was de-haploidised with purge_haplotigs v.1.0.4^70^ setting cut-offs at 35, 85 and 70x coverages to reconstruct a high-quality haploid reference assembly. HiC-based chromosome scaffolding was performed as described below. Merqury v.1.1^71^ and BUSCO v.5^72^ were used to assess genome completeness and evaluate the quality of the assembly.

### Transcriptome sequencing

Fourteen samples spanning key developmental time points of *O. fusiformis* life cycle, including active oocyte, zygote, 2-cell, 4-cell, and 8-cell stages, 3 hours post-fertilisation (hpf), 4 hpf, coeloblastula (5 hpf), gastrula (9 hpf), axial elongation (13 hpf), early larva (18 hpf), mitraria larva (27 hpf), pre-metamorphic competent larva (3 weeks post-fertilisation, wpf) and post-metamorphic juvenile were collected in duplicates (except for the latter), flash frozen in liquid nitrogen and stored at −80 °C for total RNA extraction. Samples within replicates were paired, with each one containing ∼300 embryos or ∼150 larvae coming from the same *in vitro* fertilisation. Nine further samples from adult tissues and body regions (blood vessel, body wall, midgut, prostomium, head, ovary, retractor muscle, tail, and testes) were also collected as described above. Likewise, further five samples spanning post-cleavage time points of *C. teleta*, including 64 cells and gastrula stages, and stage 4tt, stage 5, and stage 7 larval stages, were also collected in duplicates. Total RNA was isolated with the Monarch Total RNA Miniprep Kit (New England Biolabs, NEB) following supplier’s recommendations. Total RNA samples from developmental stages from both *O. fusiformis* and *C. teleta* were used to prep strand-specific mRNA Illumina libraries that were sequenced at the Oxford Genomics Centre (University of Oxford, UK) over three lanes of an Illumina NovaSeq6000 system in 2 × 150 bases mode. Adult tissue samples were sequenced at BGI on a BGISeq-500 platform in 2 × 100 bases mode. All samples were sequenced to a depth of ∼50 M reads (Supplementary Tables 13, 16).

### Annotation of repeats and transposable elements (TEs)

RepeatModeler v.2.0.1^73^ and RepBase were used to construct a *de novo* repeat library for *O. fusiformis*, which was then filtered for *bona fide* genes using the predicted proteome of *C. teleta*. Briefly, we used DIAMOND v.0.9.22^74^ with an *e*-value cut-off of 1e-10 to identify sequences in the *de novo* repeat library with significant similarity to protein coding genes in *C. teleta* that are not transposable elements. Sequences with a significant hit were manually inspected to verify they were not transposable elements and if so, they were manually removed from the *de novo* repeat library. The filtered consensus repeat predictions were then used to annotate the genome assembly of *O. fusiformis* with RepeatMasker “open-4.0”. We next used LTR_finder v.1.07^75^, a structural search algorithm, to identify and annotate Long Tandem Repeats (LTR). Finally, we generated a consensus set of repeats by merging RepeatMasker and LTR_finder predictions with RepeatCraft^76^, using default parameters but a maximum LTR size of 25 kb (as derived from the LTR_finder annotation). The general feature format (gff) and fasta files with the annotation of TEs and repeats are available in the GitHub repository (see Data Availability section).

### Gene prediction and functional annotation

We used SAMtools v.1.9^77^ and the annotation of repeats to soft-mask *O. fusiformis* genome assembly before gene prediction. We then mapped all embryonic and adult transcriptomes and a publicly available dataset^78^ (SRR1222288) with STAR v. 2.5.3a^79^ after removing low-quality read pairs and read pairs containing Illumina sequencing adapters with trimmomatic v.0.39^80^. StringTie v.1.3.6^81^ was used to convert STAR alignments into gene transfer format (GTF) files and Portcullis v.1.1.2^82^ to generate a curated set of splice junctions. Additionally, we generated *de novo* transcriptome assemblies for all samples with Trinity v.2.5.1^83^ with default parameters, which were thereafter mapped to the soft-masked assembly with GMAP v.2020-04-08^84^. We then ran the default Mikado v.2.1 pipeline^85^ to merge all transcriptomic evidence and reliable splice junctions into a single set of best-supported transcripts and gene models. From this merged dataset, we filtered full-length, non-redundant transcripts with a BLAST hit on at least 50 % of their length and at least two exons to obtain a gene set that we used to train Augustus v.3.2.3^86^. Simultaneously, we used the Mikado gene annotation and Portcullis splice junctions to generate confident sets of exon and intron hints, respectively. We also ran Exonerate v.2.4.0^87^ to generate spliced alignments of the proteome of *C. teleta* proteome on *O. fusiformis* soft-masked genome assembly to obtain further gene hints. We then merged all exon and intron hints into a single dataset which we passed to Augustus v.3.2.3^86^ for *ab initio* gene prediction. Finally, PASA v.2.3.3^88^ was used to combine RNA-seq and *ab initio* gene models into a final gene set, from which spurious predictions with in-frame STOP codons (228 gene models), predictions that overlapped with repeats (5,779 gene models) and that had high similarity to transposable elements in the RepeatPeps.lib database (2,450 models) were removed. This filtered gene set includes 26,966 genes, encompassing 31,903 different transcripts. To assess the completeness of this annotation, we ran BUSCO v.5^72^ in proteome mode, resulting in 97.7 % of the core genes present. Moreover, 31,678 out of the 31,903 (99.29%) of the filtered transcripts are supported by RNA-seq data and 80.69% of the transcripts have a significant BLAST match (*e*-value cut-off < 0.001) to a previously annotated annelid gene (database containing non-redundant proteomes of the high-quality annelid genomes of *C. teleta, D. gyrociliatus, E. andreii, L. luymesi, P. echinospica, R. pachyptila* and *S. benedictii*). A similar functional annotation approach was followed to re-annotate the genome of *C. teleta* with the new RNA-seq data, using as starting assembly the soft masked version available at Ensembl Metazoa. This resulted in 41,221 transcripts, 39,814 of which have RNA-seq support (96.59%). Additionally, 80.47% of the transcripts have a significant BLAST match (*e*-value cut-off < 0.001) to other well-annotated annelid genomes (see above).

Protein homologies for the filtered transcripts of *O. fusiformis* and *C. teleta* were annotated with BLAST v.2.2.31+^89^ on the UniProt/SwissProt database provided with Trinotate v.3.0^90^. We used HMMER v.2.3.2^91^ to identify protein domains using Trinotate’s PFAM-A database and signalP v.4.1^92^ to predict signal peptides. These functional annotations were integrated into a Trinotate database, which retrieved Gene Onthology (GO), eggNOG and KEGG terms for each transcript. In addition, we ran PANTHER HMM scoring tool to assign a PantherDB^93^ orthology ID to each transcript. In total, we retrieved a functional annotation for 22,516 transcripts (63.86 %). Functional annotation reports are provided in the GitHub repository (see Data Availability section).

### Chromosome-scale scaffolding

Sperm from a single *O. fusiformis* worm and an entire sexually mature male were used as input material to construct two Omni-C Dovetail libraries following manufacturer’s recommendations for marine invertebrates. These libraries were sequenced in an Illumina NovaSeq6000 at the Okinawa Institute of Science and Technology (Okinawa, Japan) to a depth of 229 and 247 million reads. HiC reads were processed using the Juicer pipeline r.e0d1bb7^94^ to generate a list of curated contracts (‘merged no dups’) that was subsequently employed to scaffold the assembly using 3d-dna v.180419^95^. The resulting assembly and contact map were visually inspected and curated using Juicebox v.1.11.08^94^ and adjustments submitted for a subsequent run of optimisation using 3d-dna. Finally, repeats and TEs were re-annotated in this chromosome scale assembly as described above, and the annotation obtained for the PacBio-based assembly was lifted over with Liftoff v.1.6.1^96^. All gene models but two were successfully re-annotated in the chromosome-scale assembly.

### Gene family evolution analyses

We used the AGAT suite of scripts to generate non-redundant proteomes with only the longest isoform for a set of 21 metazoan proteomes (Supplementary Table 2). To reconstruct gene families, we used OrthoFinder v.2.2.7^97^ using MMSeqs2^98^ to calculate sequence similarity scores and an inflation value of 2. OrthoFinder gene families were parsed and mapped onto a reference species phylogeny to infer gene family gains and losses at different nodes and tips using the ETE 3 library^99^, as well as to estimate the node of origin for each gene family. Gene expansions were computed for each species using a hypergeometric test against the median gene number per species for a given family employing previously published code^34^ (Supplementary Tables 3–7). Principal component analysis was performed on the orthogroups matrix by metazoan lineage, given that orthogroups were present in at least three of the 22 analysed species, to eliminate taxonomically restricted genes. All single copy ortholog files derived from this analysis employed throughout the study are available in the GitHub repository (see Data Availability section).

### Macrosynteny analyses

Single copy orthologues obtained using the mutual best hit (MBH) approach generated using MMseqs2^98^ using the annotations of *Branchiostoma floridae*^100^, *Pecten maximus*^101^, *Streblospio benedictii*^102^, and *Lineus longissimus*^103,104^ were used to generate Oxford synteny plots comparing sequentially indexed orthologue positions. Plotting order was determined by hierarchical clustering of the shared orthologue content using the complete linkage method as originally proposed. Comparison of the karyotype of all four species was performed using the Rideogram package by colouring pairwise orthologues according to the ALG assignment in comparisons with *P. maximus* and *B. floridae*.

### Evolutionary analysis of chordin in annelids

The identification of *chordin* (*chrd*) and *chordin-like* (*chrdl*) genes in *O. fusiformis* was based on the genome functional annotation (see above). To mine *chrd* orthologues, 81 annelid transcriptomic datasets were downloaded from SRA (Supplementary Table 8) and assembled with Trinity v.2.5.1^83^ to create BLAST local nucleotide databases. We also created a nucleotide database for *C. teleta* using its annotated genome^105^ (ENA accession number GCA_000328365.1). Human and *O. fusiformis* CHRD proteins were used as queries to find *chrd* orthologues following the MBH approach (*e*-value≤10^−3^), obtaining 103 unique candidate *chrd* transcripts that were then translated (Supplementary Table 9). A single candidate CHRD protein for *Themiste lageniformis* (unpublished data, provided by Michael J Boyle) was included *ad hoc* at this step. In addition, 15 curated CHRD and CHRDL protein sequences (and an outgroup) were fetched from various sources (Supplementary Table 10) and aligned together with *O. fusiformis* CHRD and CHRDL sequences in MAFFT v.7^106^ with the G-INS-I iterative refinement method and default scoring parameters. From this mother alignment further daughter alignments were obtained using “mafft --addfragments”^107^, the accurate “--multipair” method, and default scoring parameters. For orthology assignment, two phylogenetic analyses were performed on selected candidate sequences, which included the longest isoform for each species-gene combination, given that it included a 10-residue or longer properly aligned fragment in either the CHRD domains or the von Willebrand factor type C (VWFC) domains. vWFC and CHRD domains were trimmed and concatenated using domain boundaries defined by ProSITE domain annotation for the human chordin precursor protein (UniProt: Q9H2×0). Either all domains or the VWFC domains only were used for phylogenetic inference (Extended Data Figure 2c, d, Supplementary Tables 11, 12) with a WAG amino acid replacement matrix^108^ to account for transition rates, the FreeRate heterogeneity model (R4)^109^ to describe sites evolution rates, and an optimization of amino acid frequencies using maximum likelihood (ML) using IQ-TREE v.2.0.3^110^. 1,000 ultrafast bootstraps (BS)^111^ were used to extract branch support values. Bayesian reconstruction in MrBayes v.3.2.7a^112^ were also performed using the same WAG matrix but substituting the R4 model for the discrete gamma model^113^, with 4 rate categories (G4). All trees were composed in FigTree v.1.4.4. Alignment files are available in the GitHub repository (see Data Availability section).

### Gene expression profiling

We profiled gene expression dynamics from blastula to juvenile stages for *O. fusiformis*, from 64-cell to competent larva stages for *C. teleta*, from early development to female adult stages for *D. gyrociliatus*, and across the 9 adult tissues samples of *O. fusiformis*. Sequencing adaptors were removed from raw reads using trimmomatic v.0.39^80^. Cleaned reads were pseudo-aligned to the filtered gene models using kallisto v.0.46.2^114^ and genes with an expression level above an empirically defined threshold of 2 transcripts per million (TPM) were deemed expressed. For each species, the DESeq2 v.1.30.1 package^115^ was used to normalise read counts across developmental stages (Supplementary Tables 13–21) and adult tissues (Supplementary Tables 49–51) and to perform pair-wise differential gene expression analyses between consecutive developmental stages. *P-*values were adjusted using the Benjamini-Hochberg method for multiple testing correction. We defined a gene as significantly upregulated for a log_2_(fold-change) (LFC) > 1 or downregulated for a LFC < 1, given that adjusted *p*-value < 0.05. Principal component analyses were performed on the variance stabilising-transformed matrices of the normalised DESeq2 matrices. For the *O. fusiformis* adult tissues samples, genes specifically expressed (TPM > 2) in both the head and head plus two anteriormost segments samples only were classified as adult anterior genes, and those expressed in both the tail and the body wall only were classified as adult trunk and posterior genes (Supplementary Tables 52, 53). For all 3 annelid taxa, anterior, trunk, and posterior markers were defined as genes whose spatial expression pattern has been validated through *in situ* hybridisation in the literature (Supplementary Tables 54–56). TPM and DESeq2 gene expression matrices of developmental and adult tissue samples are also available in the GitHub repository (see Data Availability section).

### Gene clustering and co-expression network analyses

Transcripts were clustered according to their normalised DESeq2 expression dynamics through soft *k-*means clustering (or soft clustering) using the mfuzz v.2.52 package^116^ (Supplementary Tables 23–26). Out of the total number of transcripts, we discarded those which were not expressed at any developmental stage (225 out of 31,903 for *O. fusiformis*, 1,407 out of 41,221 for *C. teleta*, and 200 out of 17,388 for *D. gyrociliatus*). We then determined an optimal number of 12 clusters (*O. fusiformis* and *C. teleta*) and 9 clusters (*D. gyrociliatus*) for our datasets by applying the elbow method to the minimum centroid distance as a function of the number of clusters. For the construction of the gene co-expression networks for *O. fusiformis* and *C. teleta*, we used the WGCNA package v.1.70–3^117^. All transcripts expressed at any developmental stage were used to build a signed network with a minimum module size of 300 genes and an optimised soft-thresholding power of 16 and 8, for *O. fusiformis* and *C. teleta*, respectively. Block-wise network construction returned 15 gene modules for *O. fusiformis*, from which one module was dropped due to poor intramodular connectivity, and 19 gene modules for *C. teleta* (Supplementary Tables 23, 24). The remaining 14 gene modules of *O. fusiformis* (A–N) and 19 gene modules of *C. teleta* (A– O, W–Z) were labelled with distinct colours with unassigned genes labelled in grey. Random subsets consisting of the nodes and edges of 30 % of the transcripts were fed to Cytoscape v.3.8.2^118^ for network visualisation. Module eigengenes were chosen to summarise the gene expression profiles of gene modules. Gene ontology (GO) enrichment analysis of each gene cluster and gene module was performed using the topGO v.2.44 package. We performed a Fisher’s exact test and listed the top 30 (soft *k*-means clusters) or top 15 (WGCNA modules) significantly enriched GO terms of the class biological process (Supplementary Tables 27–31). To ease visualisation, all 486 non-redundant enriched GO terms from the 33 soft *k-* means clusters from all 3 species were clustered through *k*-means clustering by semantic similarity using the simplifyEnrichment v.1.2.0 package^119^ (Supplementary Figures 7, 8). Full network nodes and edges files and the random 30 % network subset files are available in the GitHub repository (see Data availability section).

### Transcription factor repertoire analysis

We selected a custom set of 36 transcription factor classes from all 9 transcription factor superclasses from the TFClass database^120^. Transcripts in *O. fusiformis, C. teleta*, and *D. gyrociliatus* were deemed transcription factors and classified into one or more of the 36 classes if they were a match for any of the corresponding PANTHER identifiers (Supplementary Tables 32–33, Supplementary Figure 3). Over- and underrepresentation of the different transcription factor classes in the gene expression clusters was tested through pair-wise two-sided Fisher’s exact tests, for which we then adjusted the *p-*values using the Benjamini-Hochberg correction for multiple testing.

### Orthogroup overlap analysis

We performed pair-wise comparisons between each possible combination of soft *k*-means clusters of all 3 annelid taxa. The numbers of overlapped orthogroups between either the full clusters or the transcription factors belonging to each cluster only were subjected to upper-tail hypergeometric tests. *P*-values were then adjusted using the Benjamini-Hochberg method for multiple testing correction. For the simplified analyses by quadrants, clusters were classed as early/pre-larval (*O. fusiformis*: 1–6; *C. teleta*: 1–5; *D. gyrociliatus*: 1–3) or late/pre-larval (*O. fusiformis*: 8–12; *C. teleta*: 7–12; *D. gyrociliatus*: 5–7), thus rendering 4 different quadrants for each species pair-wise comparison: early_species A_ –early_species B_, early_species A_ – late_species B_, late_species A_ –early_species B_, and late_species A_ –late_species B_. Clusters corresponding to female adult expression in *D. gyrociliatus* (8 and 9) were discarded for comparison purposes. Relative similarity (*RS*) for each of the four quadrants was computed as the following ratio:

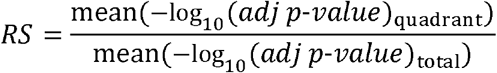

Values above 1 indicate a higher orthogroup overlap than average, whereas values below 1 represent a lower overlap than average. For genes under heterochronic shifts – i.e., with distinct temporal expression dynamics – between indirect and direct development, a gene set was constructed with the genes with a single copy ortholog in both *O. fusiformis* and *C. teleta* whose expression was shifted from post-larval clusters (*O. fusiformis*: 7–12; *C. teleta*: 8–12) to early clusters 2 and 3 in *D. gyrociliatus* (see Fig. 2f). For the characterisation of genes under heterochronic shifts between planktotrophic and lecithotrophic larvae, two gene sets were generated with the genes with early_*O. fusiformis*_–late_*C. teleta*_ and late_*O. fusiformis*_–early_*C. teleta*_ dynamics, as described above. Gene ontology (GO) enrichment analysis of both gene sets was performed using the topGO v.2.44 package. We performed a Fisher’s exact test and listed the top 15 significantly enriched GO terms of the class biological process (Supplementary Table 40). BlastKOALA^121^ server was used to assign a KEGG orthology number to one-to-one orthologs showing heterochronic sifts and KEGG mapper^122^ to analyse the annotations (Supplementary Tables 41, 42).

### Pathway analyses

Human genes involved in the animal autophagy pathway (map04140) were obtained from the KEGG pathway database^123^. *D. melanogaster* and *Saccharomyces cerevisiae* genes involved in the chitin synthesis pathway were fetched from FlyBase^124^ and SGD^125^ based on the enzyme nomenclature (EC) numbers of the pathway enzymatic activities^126^. Orthology in *O. fusiformis* and *C. teleta* for the autophagy pathway genes was determined from the single copy ortholog sets to the human genes, where one for both species existed (Supplementary Tables 43, 44). For the chitin synthesis pathway, and due to the high number of paralogs and expansions/losses of enzymatic activities of the chitin synthesis pathway, orthology was inferred from PANTHER family/subfamily identifiers to the corresponding enzymatic activities (Supplementary Tables 45, 46). We then used this orthology to reconstruct the chitin synthesis pathway in annelids. Timing across both species and the presence or lack thereof of heterochronic shifts between *O. fusiformis* and *C. teleta* was determined as described above.

### Hox genes orthology assignment

129 curated Hox sequences were retrieved from various databases (Supplementary Table 47) and aligned with *O. fusiformis* Hox proteins with MAFFT v.7 in automatic mode. Poorly aligned regions were removed with gBlocks v.0.91b^127^ yielding the final alignments. Maximum likelihood trees were constructed using RAxML v.8.2.11.9^128^ with an LG substitution matrix^129^ and 1,000 ultrafast BS. All trees were composed in FigTree v.1.4.4. Alignment files are available in the GitHub repository (see Data Availability section).

### Whole mount in situ hybridisation and immunohistochemistry

Fragments of *chordin* and *Hox* genes were isolated as previously described^27^ using gene-specific oligonucleotides and a T7 adaptor. Riboprobes were synthesise with the T7 MEGAscript kit (ThermoFisher, AM1334) and stored at a concentration of 50 ng/μl in hybridisation buffer at −20 ºC. Whole mount *in situ* hybridisation in embryonic, larval, and juvenile stages were conducted as described elsewhere^27,29^. Antibody staining in larval stages of *O. fusiformis, Magelona* spp. and *C. teleta* was carried out as previously described^26,130^. DIC images of the colorimetric *in situs* were obtained with a Leica 560 DMRA2 upright microscope equipped with an Infinity5 camera (Lumenera). Fluorescently stained samples were scanned with a Nikon CSU-W1 Spinning Disk Confocal.

### Assay for Transposase-Accessible Chromatin using sequencing (ATAC-seq)

We performed two replicates of ATAC-seq from samples containing ∼50,000 cells at the blastula (∼900 embryos), gastrula (∼500), elongation (∼300), mitraria larva (∼150 larvae) and competent larva (∼40) stages for *O. fusiformis*, and the 64-cells stage (∼500 embryos), gastrula (∼200), stage 4tt larva (∼120 larvae), stage 5 larva (∼90) and stage 8 larva (∼50) for *C. teleta* following the omniATAC protocol^131^, but gently homogenising the samples with a pestle in lysis buffer and incubating them on ice for 3 min. Tagmentation was performed for 30 min at 37°C with an in-house purified Tn5 enzyme^132^. After DNA clean-up, ATAC-seq libraries were amplified as previously described. Primers used for both PCR and qPCR are listed in Supplementary Tables 57 and 59. Amplified libraries were purified using ClentMag PCR Clean Up Beads as indicated by the supplier and quantified and quality checked on a Qubit 4 Fluorometer (Thermo-Fisher) and an Agilent 2200 TapeStation system before pooling at equal molecular weight. Sequencing was performed on an Illumina HiSeq4000 platform in 2 × 75 bases mode at the Oxford Genomics Centre (University of Oxford, United Kingdom) (blastula, elongation and mitraria larva stages, and one replicate of the gastrula sample of *O. fusiformis*, as well as the 64 cells, gastrula, and stage 4tt larva stages of *C. teleta*) and on an Illumina NovoSeq6000 in 2 × 150 bases mode at Novogene (Cambridge, United Kingdom) (one replicate of gastrula and the two replicates of competent larva stages of *O. fusiformis* and the two replicates of stage 5 and stage 8 larva of *C. teleta*).

### Chromatin accessibility profiling

We used cutadapt v.2.5^133^ to remove sequencing adaptors and trim reads from libraries sequenced in 2 × 150 bases mode to 75 bases reads. Quality filtered reads were mapped using NextGenMap v.0.5.5^134^ in paired-end mode, duplicates were removed using samtools v.1.9^135^ and mapped reads were shifted using deepTools v.3.4.3^136^ (Supplementary Tables 58, 60). Fragment size distribution was estimated from resulting BAM files and transcription start site (TSS) enrichment analysis was computed using computeMatrix and plotHeatmap commands in deepTools v.3.4.3. Peak calling was done with MACS2 v.2.2.7.1^137,138^ (-f BAMPE --min- length 100 --max-gap 75 and -q 0.01). Reproducible peaks were identified by irreproducible discovery rates (IDR) (IDR < 0.05) v.2.0.4. at each developmental stage. Peaks from repetitive regions were filtered with BEDtools v.2.28.0^139^ at each developmental stage. Next, we used DiffBind v.3.0.14^140^ to generate a final consensus peak set of 63,732 peaks in *O. fusiformis* and 46,409 peaks in *C. teleta*, which were normalised using DESeq2 method. Peak clustering according to accessibility dynamics was performed as described above for RNA-seq, using the same number of 12 clusters to make both profiling techniques comparable. Principal component analysis and differential accessibility analyses between consecutive developmental stages were also performed as described above. An LFC > 0 and a LFC < 0 indicates whether a peak opens or closes, respectively, given that the adjusted *p*-value < 0.05. Stage-specific and constitutive peaks were determined using UpSetR v.1.4.0^141^ and both the consensus peak set and the stage-specific peak sets were classified by genomic region using HOMER v.4.11^142^ and further curated. Visualisation of peak tracks and gene structures was conducted with pyGenomeTracks v.2.1^143^ and deepTools v.3.4.3^136^. To correlate chromatin accessibility and gene expression, this genomic region annotation was used to assign peaks to their closest gene (63,726 peaks were assigned to 23,025 genes in *O. fusiformis* and 44,368 peaks were assigned to 23,382 genes in *C. teleta*). Pearson correlation coefficient between chromatin accessibility and gene expression was computed individually by peak, and gene ontology (GO) enrichment analyses of the gene sets regulated by peak clusters was performed using the topGO v.2.44 package. We performed a Fisher’s exact test and listed the top 30 significantly enriched GO terms of the class biological process. To ease visualisation, all 242 non-redundant enriched GO terms were clustered through *k*-means clustering by semantic similarity using the simplifyEnrichment v.1.2.0 package^119^ (Supplementary Tables 61–71). Coverage files and peak set files are available in the GitHub repository (see Data Availability section).

### Motif identification, clustering, matching and curation

To identify transcription factor-binding motifs in chromatin accessible regions in the two species, we first used HOMER^142^ (v.4.1) to identify known and *de novo* motifs in the consensus peak sets, which yielded 456 motifs for *O. fusiformis* and 364 motifs for *C. teleta* (Supplementary Tables 72, 73). We then used GimmeMotifs v.0.16.1^144^, with a 90% similarity cut-off to cluster the motifs predicted in *O. fusiformis* and *C. teleta* into 141 consensus motifs, which we matched against four motif databases to assign their putative identity (Gimme vertebrate 5.0^144^, HOMER^142^, CIS-BP^145^ and a custom JASPAR2022^146^ core motifs without plant and fungi motifs, Supplementary Fig. 22). We then used the human non-redundant TF motif database (https://resources.altius.org/∼jvierstra/projects/motif-clustering-v2.0beta/) to manually curate the annotation. After removing motifs that likely represented sequence biases, we finally obtained 95 motif archetypes for *O. fusiformis* and 91 for *C. teleta* (Supplementary Table 74), which we then used to perform motif counts in peaks (Supplementary Tables 75, 76) and motif accessibility estimation (Supplementary Tables 77, 78) with GimmeMotifs v.0.16.1^144^. Data clustering was performed with mfuzz v.2.52^116^. Over- and underrepresentation of counts of the common curated motif archetypes in the peak accessibility soft clusters (see above) was tested through pair-wise two-sided Fisher’s exact tests, for which we then adjusted the *p-*values using the Bonferroni correction for multiple testing.

### Transcription factor footprinting and Hox gene regulatory network exploration

To predict transcription factor binding, as a proxy of activity, we conducted footprinting analysis with TOBIAS^44^ v.0.12.0 during development in the 95 and 91 motif archetypes for *O. fusiformis* and *C. teleta*, respectively (Supplementary Tables 79, 80). Transcription factor binding scores (TFBS) were clustered with mfuzz v.2.52^116^. Pearson correlation coefficients of motif accessibility and TFBS were calculated by stage and by motif separately based on the 33 common, curated motif archetypes. To reconstruct potential upstream regulators and downstream effectors of the *Hox* genes, we first subset ATAC-seq peaks annotated to the *Hox* genes in the *Hox* cluster (i.e., all but *Post1*) in *O. fusiformis* and *C. teleta* and extracted the bound motifs on those peaks (Supplementary Tables 81, 82). TFBS were sum up for each motifs to obtain global dynamics, and their temporal dynamics were then clustered with mfuzz v.2.52^116^. For the downstream genes regulated by *Hox*, we obtained genes annotated to ATAC-seq peaks with a bound HOX/EVX/CDX motif at the competent stage in *O. fusiformis* and stage 4tt larva in *C. teleta* (Supplementary Tables 83, 84). One-to-one orthologs were used to identified shared targets and PANTHER IDs to obtain their functional annotation.

### Phylostratigraphy

To evaluate gene expression dynamics by phylostratum and developmental stage in all 3 annelid lineages, we used the OrthoFinder gene families and their inferred origins. We deemed all genes originating before and with the Cnidarian-Bilaterian ancestor of pre-metazoan and metazoan origin (Supplementary Tables 85–87). We then applied a quantile normalisation onto the DESeq2 normalised matrices of gene expression. The 75 % percentile of the quantile-normalised gene expression levels was used as the summarising measure of the gene expression distribution by developmental stage. Over- and underrepresentation of the different phylostrata in the gene expression clusters was tested through pair-wise two-sided Fisher’s exact tests, for which we then adjusted the *p-*values using the Bonferroni correction for multiple testing. Gene expression dynamics of novel genes and genes of pre-metazoan and metazoan origin across selected metazoan lineages (see Comparative transcriptomics section below) were also evaluated as described above.

### Comparative transcriptomics

Publicly available RNA-seq developmental time courses for the development of *Amphimedon queenslandica, Clytia hemisphaerica, Nematostella vectensis, Strongylocentrotus purpuratus, Branchiostoma lanceolatum, Danio rerio, Drosophila melanogaster, Caenorhabditis elegans, Crassostrea gigas, Dimorphilus gyrociliatus*, and two stages of *Capitella teleta* were downloaded from the SRA (Supplementary Table 88), cleaned for adaptors and low-quality reads with trimmomatic v.0.39^80^ and pseudo-aligned to their respective non-redundant genome-based gene repertoires – i.e., with a single transcript isoform, the longest, per gene model – using kallisto v.0.46.2^114^. We then performed a quantile transformation of TPM values using scikit-learn v.1.0.2^147^ and calculated the Jensen-Shannon divergence (JSD) from (i) all single copy orthologs, (ii) the set single copy transcription factor orthologs, and (iii) the set of common single copy orthologs across all lineages, either between all possible one-to-one species comparisons (i) or between all species and *O. fusiformis* (ii, iii), using the philentropy v.0.5.0 package^148^:

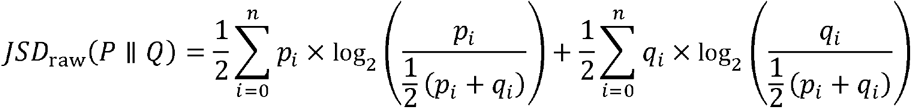

Transcriptomic divergences were calculated based on 250 bootstrap replicates, from which statistically robust mean values and standard deviations were obtained. Raw mean JSD values (*JSD*_raw_) were adjusted (*JSD*_adj_) by dividing by the number of single copy orthologs (i), single copy transcription factor orthologs (ii), or common single copy orthologs (iii) of each comparison (Supplementary Tables 89, 90), and normalised using the minimum and maximum adjusted JSD values from all one-to-one species comparisons as follows:

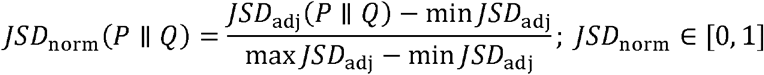

Relative JSD values were obtained equally, using minimum and maximum adjusted JSD values from each one-to-one species comparison instead. Gene-wise JSD (*gwJSD*) between five key one-to-one larval stages comparisons was computed as follows:

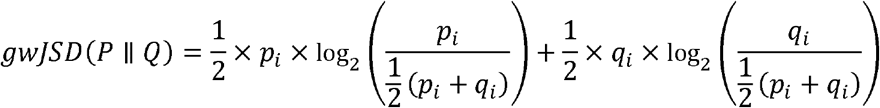

Similarity-driving genes – i.e., those with very low gwJSD – were subset as those below the threshold defined as 25 % of the point of highest probability density of the gwJSD distributions. Gene ontology (GO) enrichment analysis of the similarity-driving gene sets was performed using the topGO v.2.44 package. We performed a Fisher’s exact test and listed the top 30 significantly enriched GO terms of the class biological process (Supplementary Table 91). To ease visualisation, all 51 non-redundant enriched GO terms from the 5 gene sets were clustered through *k*-means clustering by semantic similarity using the simplifyEnrichment v.1.2.0 package^119^. For comparative *Hox* gene expression dynamics profiling in metazoan lineages, the same non-redundant gene expression matrices were normalised using the DESeq2 v.1.30.1 package^115^, unless *Hox* gene models where missing, in which case they were manually added *ad hoc* to the non-redundant genome-based gene repertoires (Supplementary Table 94). *Hox* gene expression profiling in *Urechis unicinctus* was performed as described for the rest of taxa but using the available reference transcriptome^149^ instead (Supplementary Table 48). All gene expression matrices are available in the GitHub repository (see Data Availability section).

## Data availability

All sequence data associated with this project are available at the European Nucleotide Archive (project PRJEB38497) and Gene Expression Omnibus (accession numbers GSE184126, GSE202283, GSE192478, GSE210813 and GSE210814). Genome assemblies, transposable element annotations, genome annotation files used for RNA-seq and ATAC-seq analyses, WGCNA nodes and edges files, alignment files used in orthology assignment, and other additional files are publicly available in https://github.com/ChemaMD/OweniaGenome.

## Code availability

All code used in this study is available in https://github.com/ChemaMD/OweniaGenome.

## Acknowledgments

We thank Alex de Mendoza and Daria Gavriouchkina for their support and valuable comments on the manuscript, as well as the staff at Station Biologique de Roscoff for their help with collections and animal supplies and the Oxford Genomics Centre at the Wellcome Centre for Human Genetics (funded by Wellcome Trust grant reference 203141/Z/16/Z) for the generation and initial processing of RNA-seq and ATAC-seq sequencing data. We also thank Michael J Boyle for providing the *chordin* sequence for *Themiste lageniformis*, Joseph Deane for initial help with *Hox* gene characterisation in *O. fusiformis*, and the core technical staff at the Department of Biology at Queen Mary University of London for their constant support. This work used computing resources from Queen Mary University of London’s Apocrita HPC facilities. This work was funded by the Horizon 2020 Framework Programme to JMM-D (European Research Council Starting Grant action number 801669) and AH (European Research Council Consolidator Grant action number 648861) and a Royal Society University Research Fellowship (URF\R1\191161) and a Japan Society for the Promotion of Science Kakenhi grant (JP 19K06620) to FM.

## Author Contributions

JMM-D, FM, YL and FMM-Z conceived and designed the study; YL collected RNA-seq samples for *O. fusiformis* and *C. teleta*, performed ATAC-seq experiments and contributed to all data analyses; FMM-Z performed *chordin* orthology studies and contributed to all data analyses; KG conducted *in situ* hybridisation analyses of *Hox* genes; AC-B collected RNA-seq samples for *C. teleta*, performed immunostainings on larvae and gene expression analyses of *chordin*; BED and RDD contributed to computational analyses; YT performed OMNI-C libraries; GM performed repeat annotations and analyses; OS identified and performed *in silico* analyses of *Hox* genes; MT performed genomic extractions and optical mapping; KM collected *Magelona* spp.; AH and NML contributed to sequencing efforts; FM and JMM-D assembled and annotated the genome and contributed to data analyses; YL, FMM-Z and JMM-D drafted the manuscript and all authors critically read and commented on the manuscript.

## Competing Interests

The authors declare no competing interests.

## Extended Data Figure Legends

**Extended Data Figure 1.**
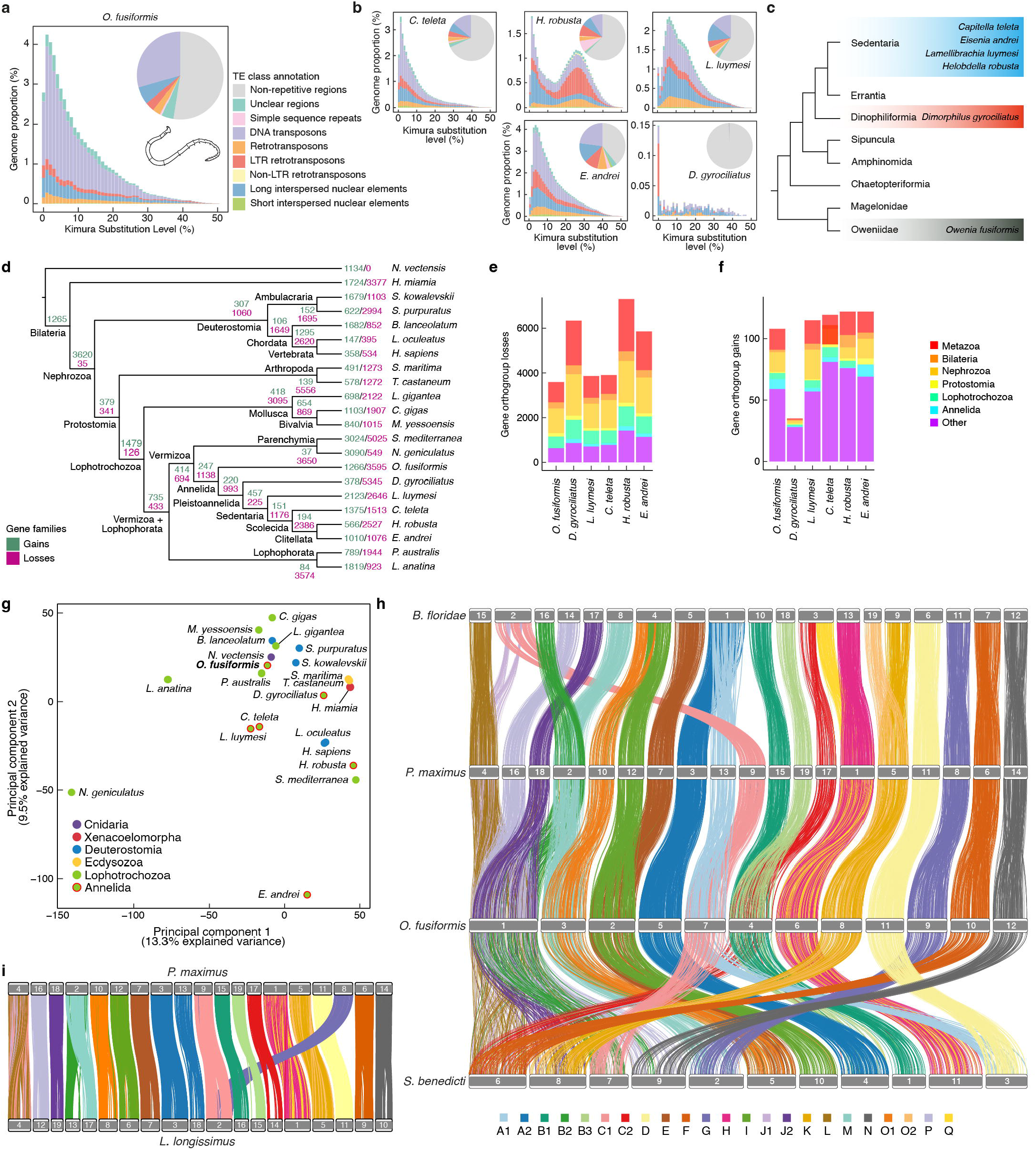
The genome of *Owenia fusiformis* is conservatively evolving. **a, b**, Pie charts of the transposable element content and Kimura substitution plots of transposable element divergence for *O. fusiformis* and other selected annelid species belonging to different annelid clades as depicted in **c**. Unlike *H. robusta* and *L. luymesi*, which show bursts of transposable elements, *O. fusiformis* shows more steady rates of expansion. **d**, Gene family evolution analysis across 22 metazoan lineages under a consensus tree topology. Gains are shown in green, losses in violet. Gene family losses in *O. fusiformis* are like those of slow-evolving lineages. **e, f**, *O. fusiformis* has the lowest number of gene losses of all sampled annelids (**e**), and the least gene expansions (**f**) after the extremely compact genome of *D. gyrociliatus*. **g**, Principal component analysis from Fig. 1f, showing the full set of species. **h**, Macrosynteny analysis between *O. fusiformis*, and from top to bottom, the cephalochordate *Branchiostoma floridae*, the bivalve *Pecten maximus*, and the annelid *Streblospio benedicti. Owenia fusiformis* retains ancestral linkage groups but also exhibits annelid- and species-specific chromosomal arrangements. However, the karyotype of *O. fusiformis* is more conserved than that of the annelid *S. benedicti*. **i**, Macrosynteny analysis between the bivalve *P. maximus* and the nemertean worm *L. longissimus. Lineus longissimus* exhibits conserved ancestral bilaterian linkage groups, including three potential lophotrochozoan-specific chromosomal rearrangements (H+Q, J2+L and K+O2), plus a nemertean-specific fusion (G+C1).

**Extended Data Figure 2.**
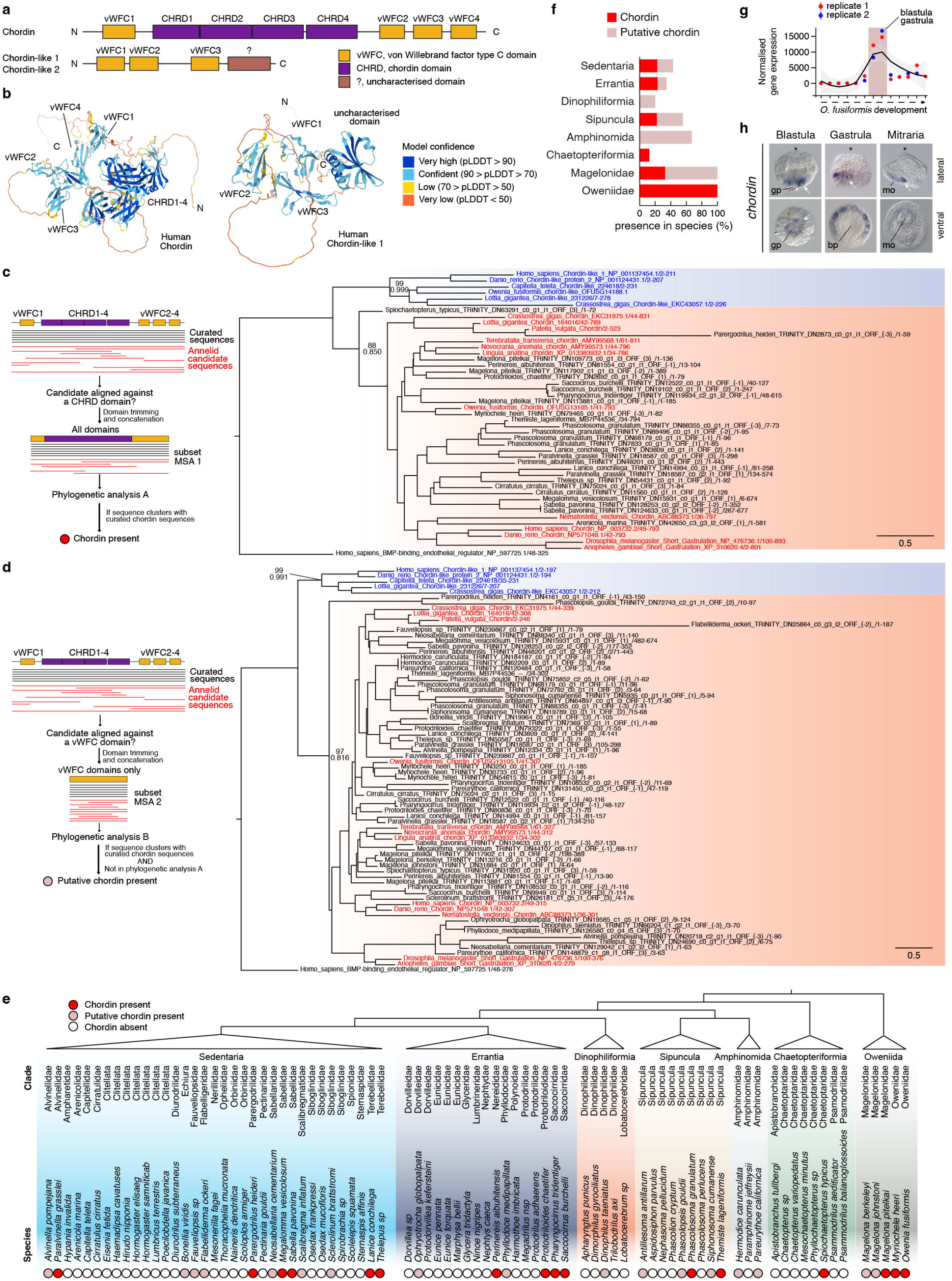
*chordin* was lost multiple times in annelids. **a**, Domain organisation of Chordin (CHRD) and Chordin-like (CHRDL1/2) proteins, as inferred from human orthologs. **b**, AlphaFold protein structure prediction for human Chordin (UniProt: Q9H2×0) and Chordin-like 1 (UniProt: Q9BU40) revealed a previously unknown and uncharacterised domain in CHRDL1 (also depicted in **a**). **c, d**, Orthology assignment of *chordin* annelid candidates. From the multiple sequence alignment, candidate annelid sequences with a 10-residue or longer fragment aligned against either the CHRD (**c**; i.e., bona fide *chordin* genes) or the vWFC domains (**d**; i.e., putative *chordin* genes) were kept for further analysis. CHRDL cluster is shaded in blue; CHRD cluster, in red. Bootstrap support values (top) and posterior probabilities (bottom) are shown at key nodes. Sequences in red and blue are curated CHRD and CHRDL sequences, respectively. **e, f**, Summary phylogenetic trees of presence/absence of *chordin* (red) or putative chordin (light brown) across Annelida. **g**, Expression levels of *chordin*, which peaks at the blastula and gastrula stages, after the specification and inductive activity of the embryonic organiser. **h**, Whole mount *in situ* hybridisation of *chordin* at the blastula (5 hours post fertilisation, hpf), gastrula (9 hpf), and mitraria larva (27 hpf) stages. Asterisks mark the animal/anterior pole. gp: gastral plate; bp: blastopore, mo: mouth.

**Extended Data Figure 3.**
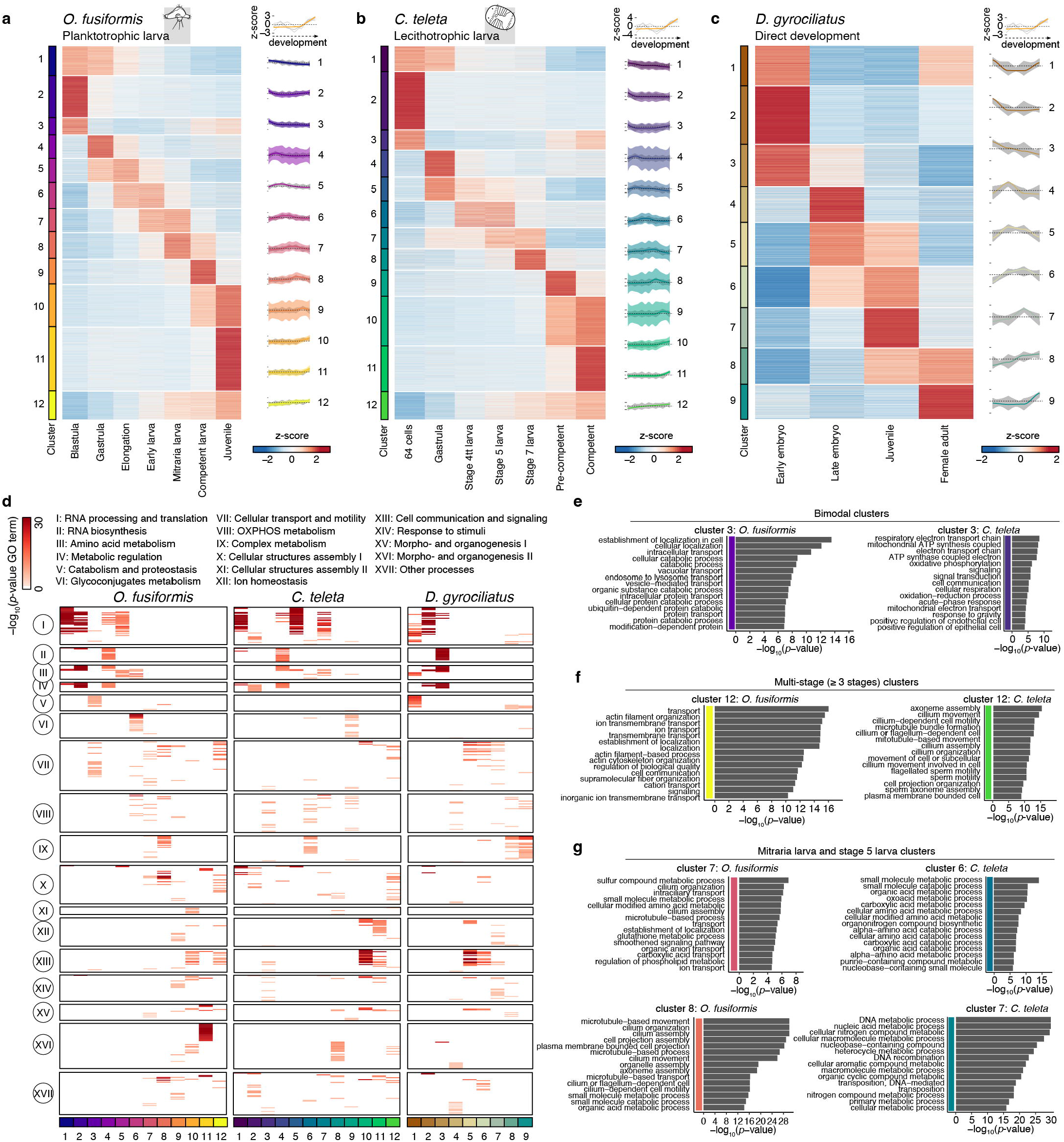
Gene expression dynamics during annelid life cycles. **a–c**, Soft *k*-means clustered heatmap of all transcripts whose expression was not null in at least one developmental stage into an optimal number of 12 clusters (*O. fusiformis*, **a**; and *C. teleta*, **b**) and 9 clusters (*D. gyrociliatus*, **c**). Soft clustering increased temporal resolution for the RNA- seq time course of *D. gyrociliatus*. On the right of each heatmap, gene-wise expression dynamics (grey lines) and locally estimated scatterplot smoothing (coloured lines) for each cluster. Coloured shaded areas represent standard error of the mean. **d**, Enrichment analysis of biological process gene ontology (GO) terms for RNA-seq clusters. Each line represents a single GO term, for which the −log^10^(*p*–value) for each RNA-seq cluster is shown in a colour coded scale. GO terms were clustered into 15 distinct clusters based on semantic similarity (see Supplementary Fig. 7, 8). Clusters are shown on the bottom of the heatmaps. **e–g**, Bar plots depicting *p*–values of the top 15 enriched GO terms in clusters with bimodal dynamics (**e**), clusters spanning three or more developmental stages (**f**) and larval clusters (**g**) for *O. fusiformis* (left) and *C. teleta* (right). For the full list of GO terms and clusters, see Supplementary Fig. 4–6.

**Extended Data Figure 4.**
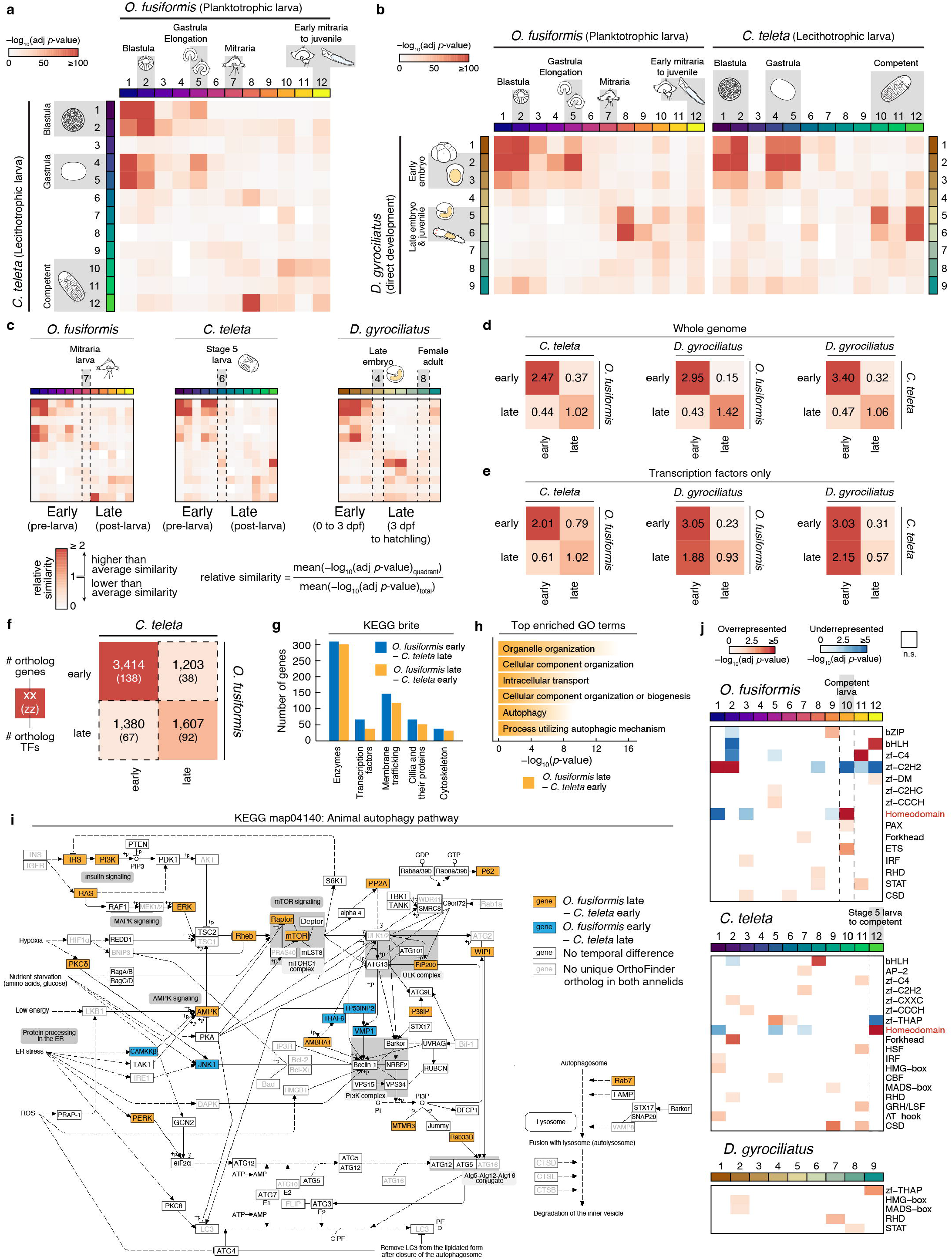
Heterochronic shifts in gene regulatory programmes between annelid life cycles. **a, b**, Similarity heatmaps showcasing the orthogroup overlap between the clusters of co-regulated genes (see Extended Data Fig. 3a–c), between the three annelids. **c**, Explanation of the orthogroup overlap analysis by quadrants. Clusters were classed as “early” (before dotted lines) or “late” (after dotted lines). Clusters of the female adult of *D. gyrociliatus* were disregarded. **d, e**, Heatmaps of relative similarity by quadrants of the orthogroup overlap analyses of the whole genomes (**d**) and transcription factors only (**e**). **f**, Single copy ortholog and transcription factor ortholog number by quadrants in the *O. fusiformis* and *C. teleta* comparison. Colour scale in **d–f** is the same as in **c. g**, KEGGbrite characterisation of the gene sets under heterochronic shifts (surrounded by dotted lines in **f**) between *O. fusiformis* and *C. teleta*. **h**, Bar plots depicting *p*-values of top biological process GO terms of genes shifted from late expression in *O. fusiformis* to early expression in *C. teleta*. Full list is available in Supplementary Fig. 12. **i**, Heterochronic shifts of genes involved in the autophagy pathway between *O. fusiformis* and *C. teleta*. 80 % of the genes under heterochronic shifts (19 of 24) are displaced from post-larval expression in *O. fusiformis* to pre-larval expression in *C. teleta*. **j**, Enrichment analysis of the number of transcription factors per class in clusters of co-transcribed genes of *O. fusiformis* (top), *C. teleta* (centre) and *D. gyrociliatus* (bottom). For each cluster and class combination, the Fisher’s exact test adjusted *p*–value is shown. Cells in red represent overrepresented classes (odds ratio, OR > 1; adjusted *p*–value < 0.05); cells in blue, underrepresented classes (OR < 1, adjusted *p*–value < 0.05). Dotted lines highlight clusters of maximal enrichment of the homeodomain class. n.s.: not significant.

**Extended Data Figure 5.**
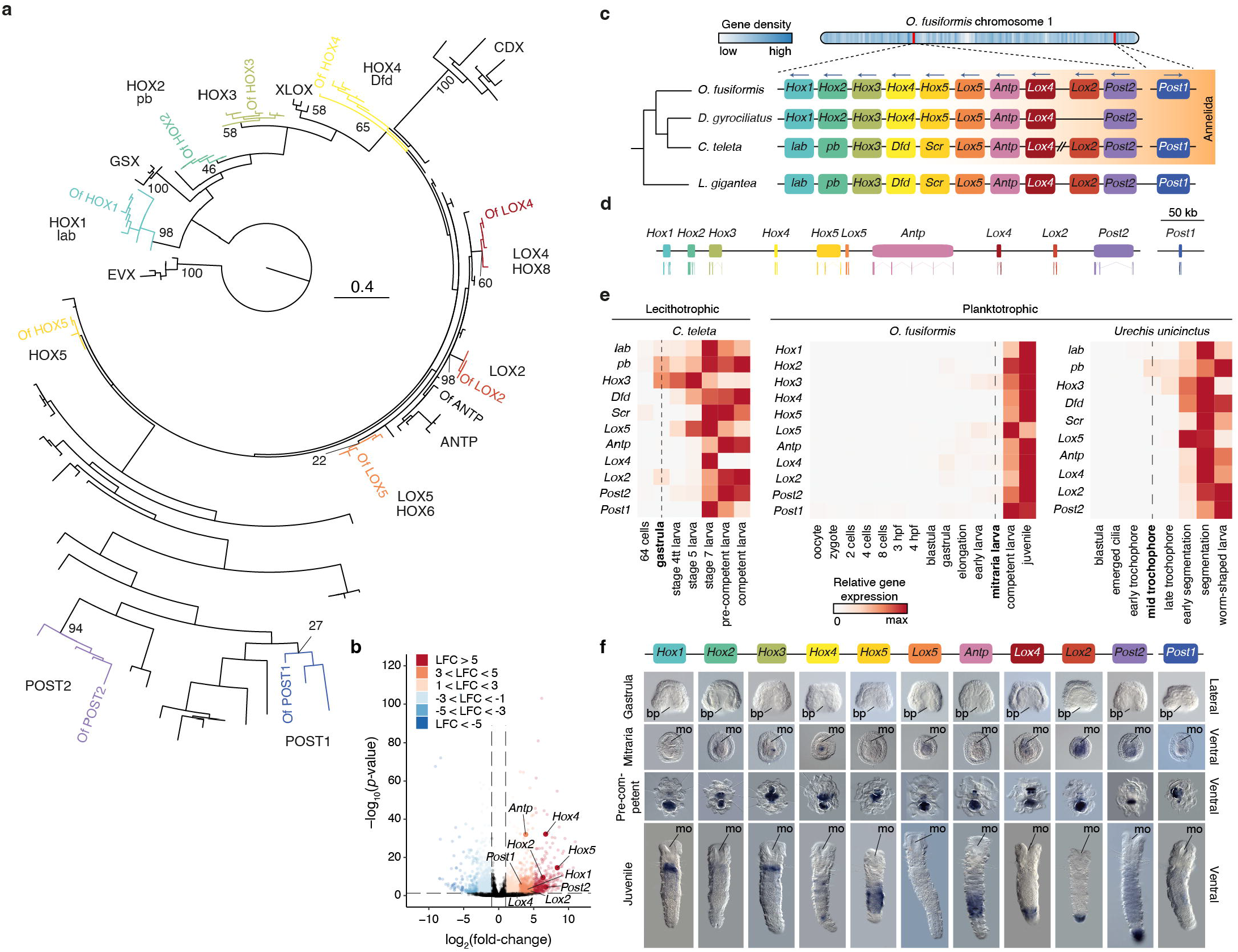
The *Hox* gene complement and expression in *O. fusiformis*. **a**, Orthology assignment of *O. fusiformis Hox* genes through maximum likelihood phylogenetic inference. Bootstrap support values are shown for major gene groups. Of: *O. fusiformis*. **b**, Volcano plot of the mitraria to competent larva transition, highlighting the marked upregulation of *Hox* genes. LFC: log2(fold-change). **c**, Chromosomal location of the *Hox* cluster and *Post1* gene in *O. fusiformis* (top) and schematic comparison of *Hox* cluster organisation in annelids and a mollusc (bottom). **d**, Schematic representation to scale of the genomic loci and intron–exon composition of *Hox* genes in *O. fusiformis*. **e**, Heatmaps of *Hox* gene expression during *C. teleta, O. fusiformis* and the echiuran annelid *Urechis unicinctus* development. In the two annelid species with planktotrophic larvae, *Hox* genes only become expressed at the larval stage (dotted vertical line) and not during embryogenesis, as observed in *C. teleta*. **f**, Whole mount *in situ* hybridisation of *Hox* genes in the gastrula (lateral views) and in the mitraria, pre-competent, and juvenile stages (ventral views). The area encircled by a dotted white line at the pre-competent stage highlights a region of probe trapping from ingested food content. bp: blastopore; mo: mouth.

**Extended Data Figure 6.**
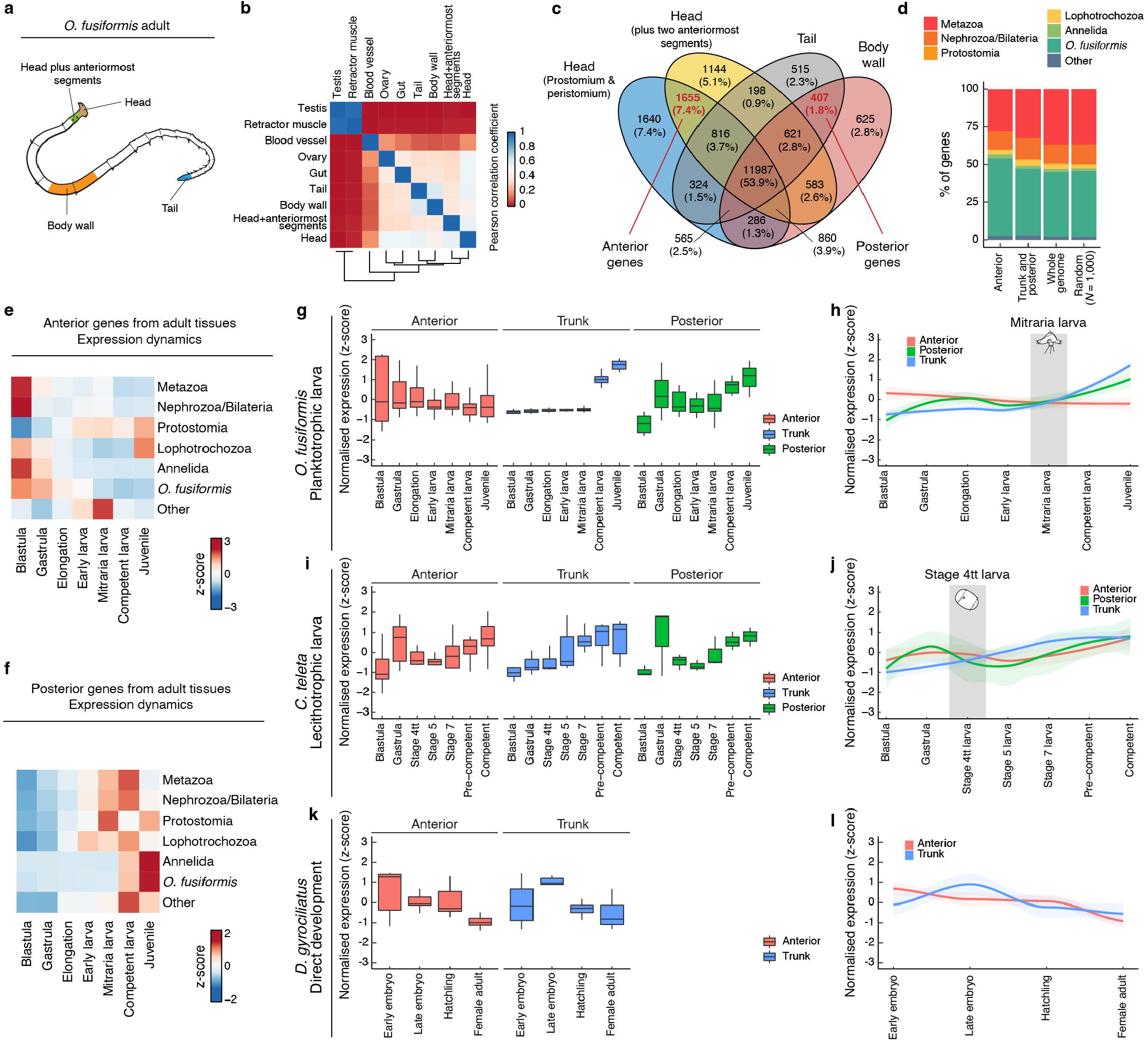
Transcriptomic profiling of adult tissues and anterior-posterior expression dynamics. **a**, Drawing of an *O. fusiformis* adult. Samples of highlighted body parts were used to define adult anterior and posterior/trunk genes. **b**, Correlation matrix of RNA-seq experiments from all nine adult tissues, calculated from a variance stabilising-transformed matrix of the normalised DESeq2 matrix. Tail and body wall samples, and head and head plus anteriormost segments samples are the most correlated samples to each other. **c**, Venn diagram showing the number of tissue-specific and shared expressed genes (TPM > 2). Gene sets highlighted with red text were defined as adult anterior, and adult trunk and posterior genes. **d**, Phylostratigraphic classification of adult anterior, and adult trunk and posterior genes, compared to the whole genome and a random subset of 1,000 genes. See Extended Data Figure 8a for phylostrata explanation. Adult genes are enriched in *O. fusiformis*-specific gene innovations. **e, f**, Expression dynamics of each phylostratum by developmental stage in the adult anterior (**e**) and adult trunk and posterior gene sets (**f**), calculated from the 75 % percentile of a quantile-normalised matrix of gene expression levels. Adult anterior genes of most phylostrata peak at the blastula, with the maximum expression of adult trunk and posterior genes of most phylostrata at post-larval stages. **g–l**, Expression dynamics of *in situ* hybridisation-validated anterior, trunk, and posterior markers throughout *O. fusiformis* (**g, h**), *C. teleta* (**i, j**), and *D. gyrociliatus* (**k, l**) development. Curves in **h, j**, and **l** are locally estimated scatterplot smoothings. Coloured shaded areas represent standard error of the mean. Key stages where expression of trunk markers is incipient are shown for both *O. fusiformis* and *C. teleta*.

**Extended Data Figure 7.**
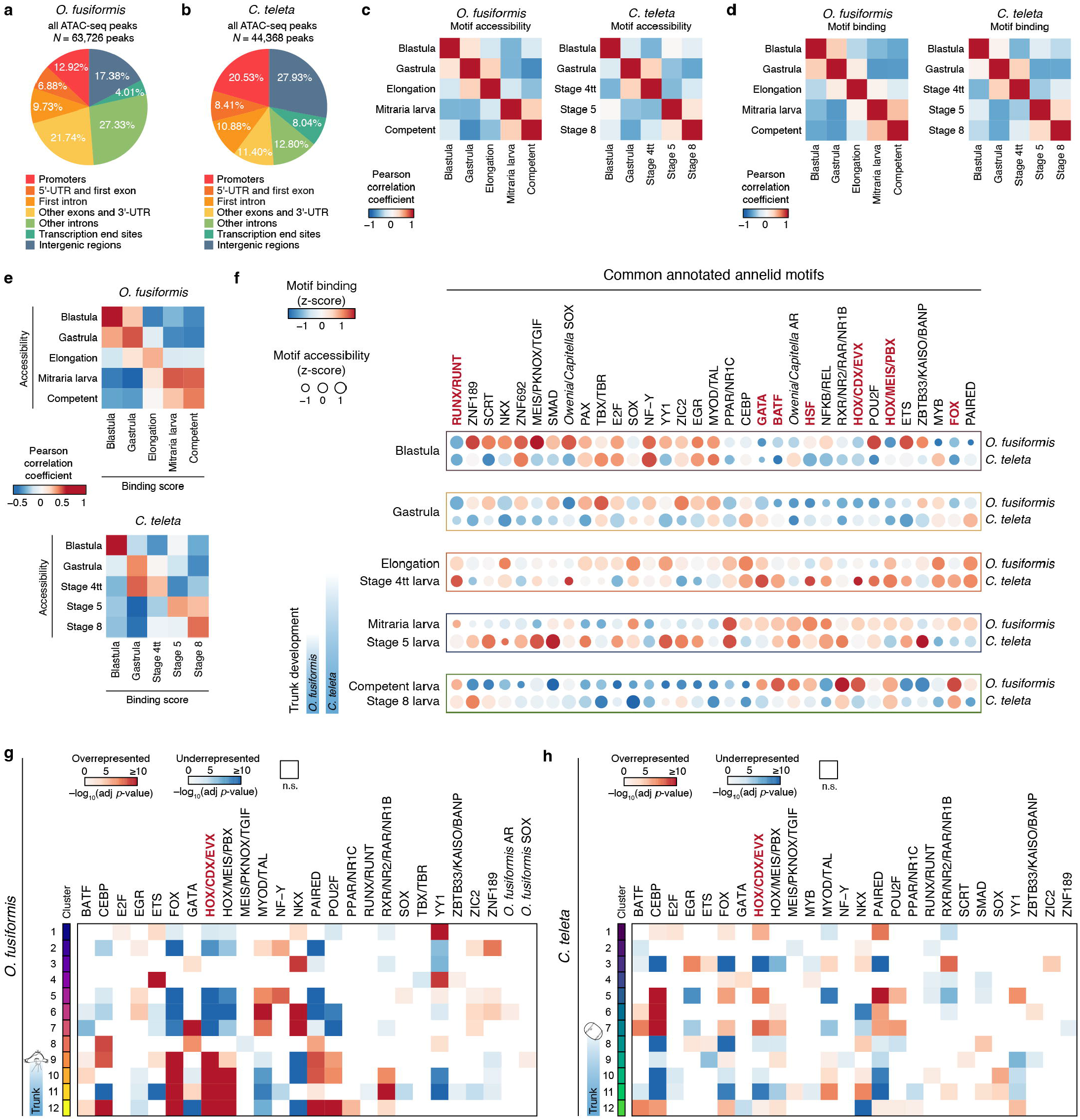
Chromatin dynamics during the development of *O. fusiformis* and *C. teleta*. **a, b**, Genomic feature annotation of the consensus ATAC-seq peak sets of *O. fusiformis* (**a**) and *C. teleta* (**b**). **c, d**, Self-correlation matrices of normalised motif accessibility (**c**) and transcription factor binding score (**d**) during *O. fusiformis* (left) and *teleta* (right) development. Both matrices demonstrate distinct chromatin regulatory dynamics at each stage of development for both species. **e**, Correlation matrices of normalised motif accessibility to transcription factor binding score during *O. fusiformis* (top) and *C. teleta* (bottom) development. Matrices demonstrate a similarity diagonal between both variables for both species. **c, d**, and **e** further validate the non-triviality of the results obtained in Fig. 4e. **f**, Heatmap of normalised motif accessibility and transcription factor binding dynamics for each of the common annotated annelid motif archetypes during *O. fusiformis* and *C. teleta* development. Colour scale denotes transcription factor binding score dynamics, bubble size represents motif accessibility dynamics, both in a z-score scale. Motif archetypes highlighted in red are representative examples of the heterochronic shifts shown in bulk in Fig. 4e. **g, h**, Enrichment analysis of the number of occurrences of the common annotated annelid motif archetypes in the peak clusters inferred through soft *k*-means clustering and shown in Fig. 4c, for *O. fusiformis* (**g**) and *C. teleta* (**h**). For each cluster and motif combination, the Bonferroni-adjusted *p*–value of the Fisher’s exact test is shown. Red cells represent significantly overrepresented lineages (odds ratio, OR > 1, adjusted *p*– value < 0.05). Blue cells denote significantly underrepresented lineages (OR < 1, adjusted *p*– value < 0.05).

**Extended Data Figure 8.**
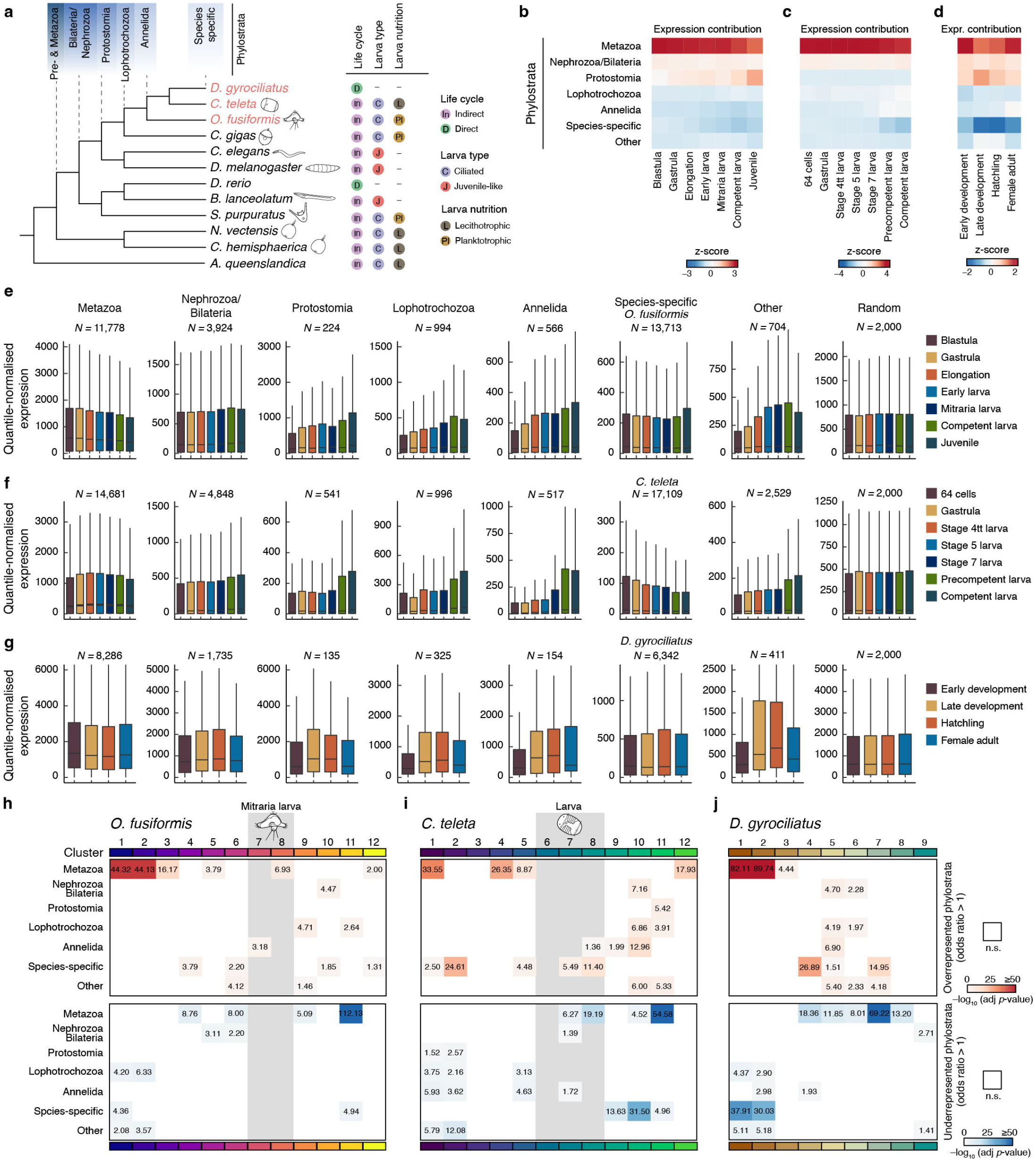
Phylostratography analyses in annelid life cycles. **a**, Cladogram of species used for comparative transcriptomics analyses, indicating on top the phylogenetic age of each phylostratum considered for phylostratigraphy analysis. For each species, the type of life cycle (direct/indirect), larval type (ciliated/juvenile-like) and larval nutritional mode (planktotrophic/lecithotrophy) are shown on the right. **b–d**, Expression contribution of each phylostratum by developmental stage in *O. fusiformis* (**b**), *C. teleta* (**c**), and *gyrociliatus* (**d**), calculated from 75% percentiles of quantile-normalised matrices of gene expression levels. Older genes are expressed at the highest levels across annelid development. **e–g**, Boxplots of quantile-normalised expression levels of genes classified by phylostratum across *O. fusiformis* (**e**), *C. teleta* (**f**), and *D. gyrociliatus* (**g**) development. A random subset of 2,000 genes is shown as a negative control. *N* denotes number of genes per phylostratum. **h–j**, Enrichment analysis of the number of genes per phylostratum in clusters of co-transcribed genes as inferred through soft *k*-means clustering and shown in Extended Data Fig. 3a –c, for *O. fusiformis* (**h**), *C. teleta* (**i**), and *D. gyrociliatus* (**j**). For each cluster and phylostratum combination, the Bonferroni-adjusted *p*–value of the Fisher’s exact test is shown. Upper tables include significantly overrepresented lineages (odds ratio, OR > 1, adjusted *p*–value < 0.05). Lower tables include significantly underrepresented lineages (OR < 1, adjusted *p*–value < 0.05). Shaded grey areas indicate clusters of genes with peak expression at the mitraria larva (*O. fusiformis*) and stage 4–7 larval stages (*C. teleta*).

**Extended Data Figure 9.**
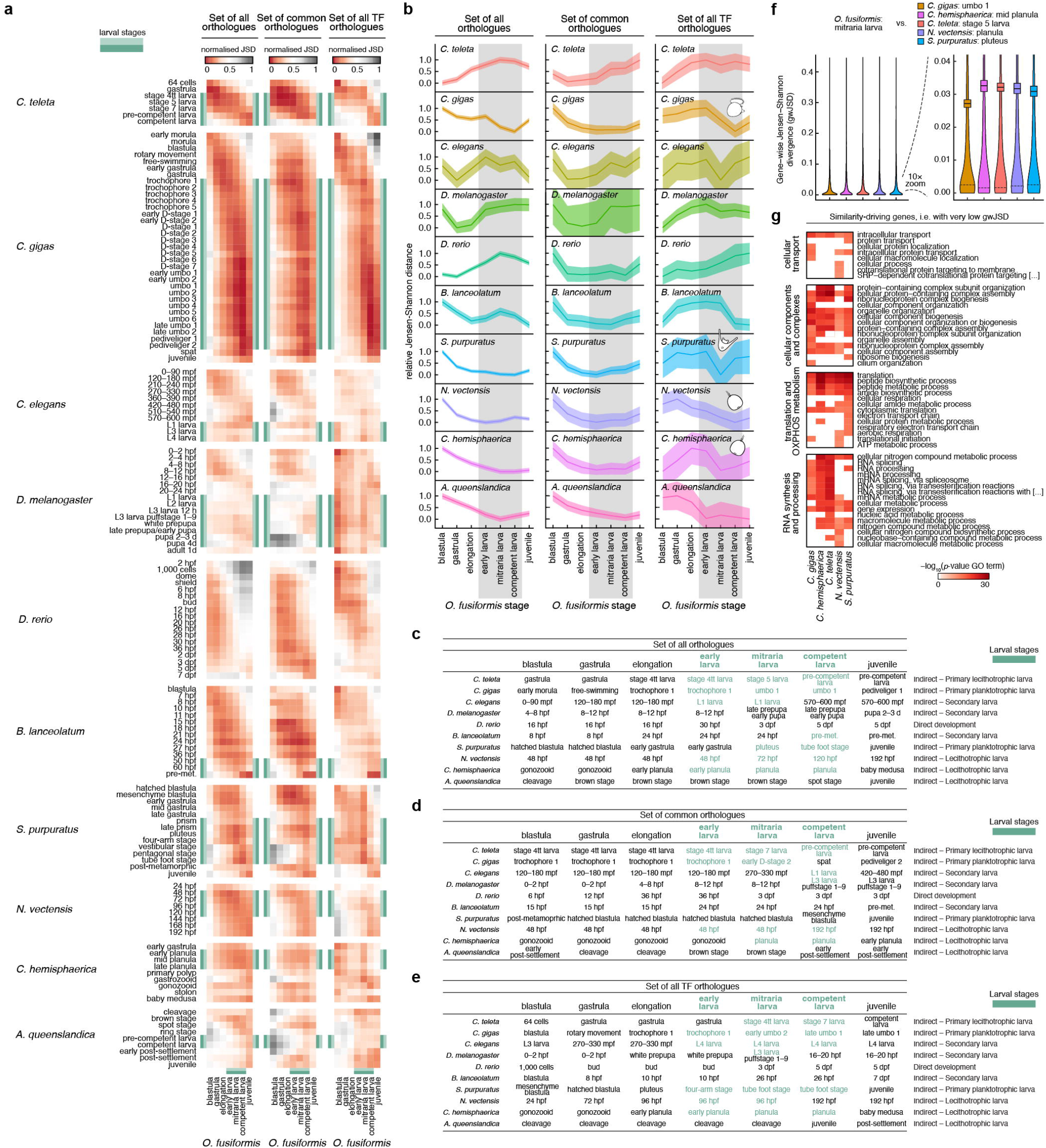
Bilaterian planktotrophic larvae and cnidarian larvae share maximal transcriptional similarity. **a**, Heatmaps of normalised transcriptomic Jensen– Shannon divergence (JSD) from pair-wise comparisons of all single copy one-to-one orthologs (left), the set of common orthologs to all species (centre), and all single copy one-to-one transcription factor orthologs (right), between *O. fusiformis* and ten other metazoan lineages with different life cycles. From top to bottom: the annelid *C. teleta*, the bivalve *C. gigas*, the nematode *C. elegans*, the insect *D. melanogaster*, the vertebrate *D. rerio*, the cephalochordate *B. lanceolatum*, the sea urchin *S. purpuratus*, the cnidarians *N. vectensis* and *C. hemisphaerica*, and the poriferan *A. queenslandica*. Larval stages are highlighted in green. **b**, Relative JSD for the datasets shown in **a**, from stages of minimal JSD to each *O. fusiformis* stage. Confidence intervals represent the standard deviation from 250 bootstrap resamplings of the ortholog sets. **c–e**, Stages of minimal JSD to each *O. fusiformis* stage, calculated from the one-to-one ortholog set (**c**), the common ortholog set (**d**), and the one-to-one transcription factor ortholog set (**e**). Larval stages are highlighted in green. **f**, Violin plots of the gene-wise Jensen Shannon divergence (gwJSD) distributions for the pair-wise comparisons of the one-to-one ortholog sets between the mitraria larva of *O. fusiformis* and the stages of minimal transcriptomic divergence as in **c**. for *C. gigas, C. hemisphaerica, C. teleta, N. vectensis*, and *S. purpuratus*. Boxes represent mean estimate and standard deviation. Dotted lines mark the point of highest probability density. Genes below ¼ of this point were subset as similarity- driving genes. **g**, Biological process GO terms enrichment of the five similarity-driving gene sets. GO terms were clustered by semantic similarity into 4 clusters. Each row represents a single GO term, for which the −log_10_(*p*–value) for each gene set is shown in a colour coded scale.

**Extended Data Figure 10.**
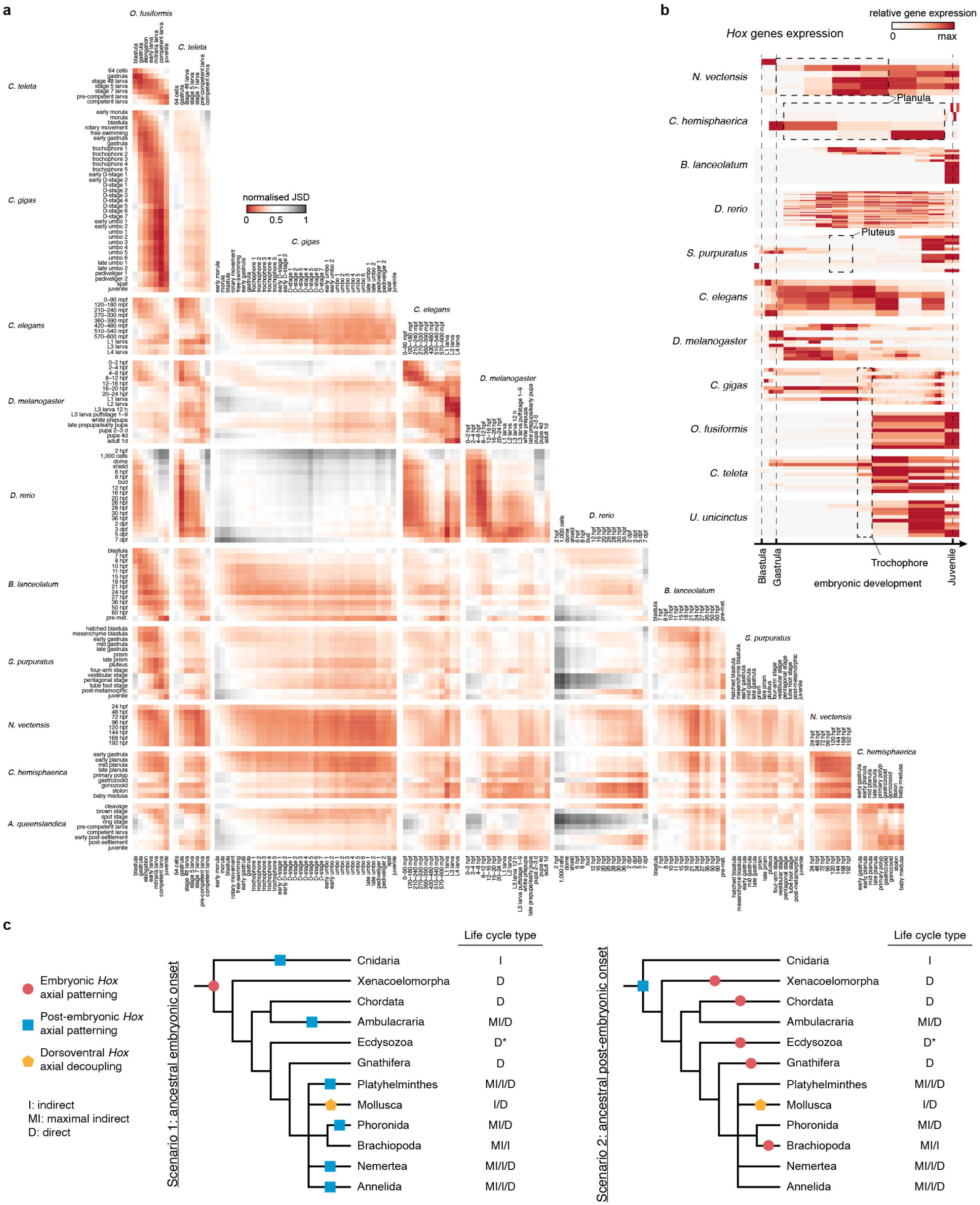
Comparative transcriptomic analysis of metazoan life cycles. **a**, Matrix of heatmaps of normalised transcriptomic Jensen–Shannon divergence (JSD) from pair-wise comparisons of all single copy one-to-one orthologs between all eleven metazoan lineages. From top to bottom and left to right: the annelids *O. fusiformis* and *C. teleta*, the bivalve *C. gigas*, the nematode *C. elegans*, the insect *D. melanogaster*, the vertebrate *D. rerio*, the cephalochordate *B. lanceolatum*, the sea urchin *S. purpuratus*, the cnidarians *N. vectensis* and *C. hemisphaerica*, and the poriferan *A. queenslandica*. **b**, Expression dynamics of *Hox* genes across the developmental RNA-seq time courses of all eleven species from **a** and the echiuran annelid *U. unicinctus*. Heatmaps were vertically aligned at the blastula, gastrula, and juvenile stages for all species. Lophotrochozoan lineages with trochophore larvae were also vertically aligned at the trochophore stage. Dotted lines encompass the larval stages of species with ciliated larvae. See Extended Data Figure 5d Supplementary Figure 32 for the fully labelled and non-deformed heatmaps. **c**, Alternative evolutionary scenarios for the deployment of *Hox* genes (as proxy for trunk patterning and assuming the staggered expression along the directive axis of cnidarians and anteroposterior axis of bilaterians is homologous, which does not necessarily imply homology of the two axes). Given our current understanding of *Hox* gene deployment in cnidarian and bilaterian taxa, a late post-embryonic *Hox* patterning ancestral to Bilateria and Cnidaria, as seen in extant lineages with maximal indirect development, is a more parsimonious scenario (on the right).

## Notes

https://github.com/ChemaMD/OweniaGenome

### Competing Interest Statement

The authors have declared no competing interest.

### Summary of Updates

Extended transcriptomic and epigenomic comparisons to support heterochronic shifts in trunk development correlate with changes in life cycle strategies.

